# The KU70-SAP domain has an overlapping function with DNA-PKcs in limiting the lateral movement of KU along DNA

**DOI:** 10.1101/2024.08.26.609806

**Authors:** Yimeng Zhu, Brian J. Lee, Shingo Fujii, Sagun Jonchhe, Hanwen Zhang, Angelina Li, Kyle J. Wang, Eli Rothenberg, Mauro Modesti, Shan Zha

**Affiliations:** Institute for Cancer Genetics, Vagelos College of Physicians and Surgeons, Columbia University, New York City, NY 10032; Department of Genome Integrity, Cancer Research Center of Marseille, CNRS UMR7258, Inserm U1068, Institut Paoli-Calmettes, Aix Marseille University, 13273 Marseille, France; Department of Biochemistry and Molecular Pharmacology, New York University School of Medicine, New York, NY 10016, USANew York University, School of Medicine; New York, NY 10016, USA; Division of Pediatric Oncology, Hematology and Stem Cell Transplantation, Department of Pediatrics, Vagelos College of Physicians & Surgeons, Columbia University, New York City, NY 10032; Department of Pathology and Cell Biology, Vagelos College of Physicians and Surgeons, Columbia University, New York City, NY 10032; Department of Immunology and Microbiology, Vagelos College of Physicians and Surgeons, Columbia University, New York City, NY 10032

**Keywords:** KU70, SAP domain, DNA-PKcs, NHEJ, V(D)J recombination, class switch recombination

## Abstract

The non-homologous end-joining (NHEJ) pathway is critical for DNA double-strand break repair and is essential for lymphocyte development and maturation. The Ku70/Ku80 heterodimer (KU) binds to DNA ends, initiating NHEJ and recruiting additional factors, including DNA-dependent protein kinase catalytic subunit (DNA-PKcs) that caps the ends and pushes KU inward. The C-terminus of Ku70 in higher eukaryotes includes a flexible linker and a SAP domain, whose physiological role remains poorly understood. To investigate this, we generated a mouse model with knock-in deletion of the SAP domain (*Ku70^ΔSAP/ΔSAP^*). *Ku70^ΔSAP^* supports KU stability and its recruitment to DNA damage sites *in vivo*. In contrast to the growth retardation and immunodeficiency seen in *Ku70^−/−^* mice, *Ku70^ΔSAP/ΔSAP^* mice show no defects in lymphocyte development and maturation. Structural modeling of KU on long dsDNA, but not dsRNA suggests that the SAP domain can bind to an adjacent major groove, where it can limit KU’s rotation and lateral movement along the dsDNA. Accordingly, in the absence of DNA-PKcs that caps the ends, Ku70^ΔSAP^ fails to support stable DNA damage-induced KU foci. In *DNA-PKcs^−/−^* mice, *Ku70^ΔSAP^* abrogates the leaky T cell development and reduces both the qualitative and quantitative aspects of residual V(D)J recombination. In the absence of DNA-PKcs, purified Ku70^ΔSAP^ has reduced affinity for DNA ends and dissociates more readily at lower concentration and accumulated as multimers at high concentration. These findings revealed a physiological role of the SAP domain in NHEJ by restricting KU rotation and lateral movement on DNA that is largely masked by DNA-PKcs.

**Highlight:** Ku70 is a conserved non-homologous end-joining (NHEJ) factor. Using genetically engineered mouse models and biochemical analyses, our study uncovered a previously unappreciated role of the C-terminal SAP domain of Ku70 in limiting the lateral movement of KU on DNA ends and ensuring end protection. The presence of DNA-PKcs partially masks this role of the SAP domain.

## Introduction

DNA double-strand breaks (DSB) represent the most severe form of DNA damage. The non-homologous end-joining (NHEJ) pathway is one of the two major DSB repair pathways in mammals(1–3). Conceptually, the mammalian NHEJ can be divided into five mostly discrete, temporally defined phases: end sensing, end protection, end tethering, end processing, and end ligation (4, 16, 21). While end-sensing is the initial event and end-ligation concludes the NHEJ, the intermediate steps, especially end tethering and end protection, might be simultaneous or interchangeable. All of the five phases requires the KU heterodimer. Currently, the mammalian NHEJ pathway includes eight known components, KU70/KU80 (KU86 for humans) heterodimer (KU), X-ray repair cross-complementing protein 4 (XRCC4), Ligase IV (LIG4), XRCC4-like factor (XLF, also called Cernunnos, gene name NHEJ1), paralog of XRCC4 and XLF (PAXX), DNA-dependent protein kinase catalytic subunit (DNA-PKcs, gene name PRKDC), and Artemis nuclease (gene name DCLRE1C)(4–6). Among them, KU70, KU80, XRCC4, LIG4, and NHEJ1 are evolutionarily conserved from yeast to humans and considered ‘core’ NHEJ factors. In contrast, DNA-PKcs and Artemis are predominantly found in vertebrates. The complete loss of DNA-PKcs and Artemis abrogates end-processing while preserving most end-ligation, despite the role of DNA-PKcs in end-tethering and end-protection.

Recent advances in single-molecule and Cryo-EM studies revealed significant molecular details and functional redundancies of the NHEJ pathway (22–27). Briefly, KU initiates NHEJ by sensing and binding to the double-stranded DNA (dsDNA) ends with nanomolar affinity, crucial for recruiting all other NHEJ factors (7,8). KU70 and KU80 share a conserved core region, including the von Willebrand factor A (vWA) domains and β-barrels, which bind together to form a stable heterodimeric ring – KU (9). The vWA domain from wings on either side, and the KU ring encircles approximately 15-20 bp of dsDNA, with KU70 located near the ends (9,10). KU binding physically blocks exonuclease-mediated end resection and helicase unwinding (10, 11), necessary for homology dependent repair, while simultaneously provides the platform to recruit other NHEJ factors for end-protection (DNA-PKcs), end-processing (DNA-PKcs, Artemis), end-tethering (DNA-PKcs, XLF, PAXX), and eventuall end-ligation (LIG4)(11,12). As such, KU binding represents a commitment to the NHEJ pathway and KU ring has to be removed for pathway switching. Specifically, DNA-PKcs binds to DNA-bound KU on the KU70 side, lifts the KU70 vWA domain, and pushes the KU ring to rotate inward (10,13–16). Notably, DNA-PKcs binds to DNA-bound KU on the KU70 side, lifting the KU70 vWA domain and rotating the KU ring inward (12–16). This action allows the N- and M-HEAT repeats of DNA-PKcs to clamp the last 10 bp near the ends, protecting the 5’ phosphate DNA ends from nuclease and ligase activity (10,13,17).

For ligation to occur, the two DNA ends must be physically brought together. Unlike homology-dependent repair mechanisms, where extensive base-pairing holds the two DNA ends together to ensure template-dependent DNA synthesis, NHEJ ligates two DNA ends with little or no homology, underscoring the importance of end-tethering. Three molecular bridges, all KU-dependent, have been identified. First, the interaction between the KU80 C-terminus and DNA-PKcs on opposite ends, and the interaction between two DNA-PKcs molecules on each end, can tether the DNA ends into a long-range synaptic complex (15,18,19). Additionally, conserved binding sites for the C-terminal tails of PAXX and XLF, located under the vWA domains of KU70 and KU80, respectively, provide two additional tethering mechanisms via the homodimers of PAXX and XLF (15,16,20–22). Finally, XLF interacts with the XRCC4 homodimer on each side of the head domain, and XRCC4 binds to LIG4, which in turn binds to KU, constitute another short-range tethering mechanism(15,21,22). The DNA-bound DNA-PKcs also recruits and activates Artemis endonuclease for end-processing (10,14). Notably, DNA-bound KU activates DNA-PKcs kinase activity, promoting the release of DNA-PKcs, which is necessary for end-ligation and transition to short-range ligation complext. Correspondingly, the presence of a kinase-dead DNA-PKcs, but not the absence of DNA-PKcs, blocks end-ligation in mice (17). Finally, KU recruits LIG4-XRCC4 to completes end-ligation (16,23).

Beyond general DSB repair, NHEJ is exclusively required for V(D)J recombination that assembles the antigen receptor gene products in developing lymphocytes. Lymphocyte specific RAG endonuclease initiates V(D)J recombination by generating two blunt signaling ends (SEs) and two hairpin coding ends (CEs). In the subsequent reaction phase, the SEs are directly and precisely joined via NHEJ to form a signaling joint (SJ). DNA-PKcs and Artemis open the hairpins at the CEs, which are then ligated to form a coding joint (CJ)(24). The CJ encodes the variable region exon of B and T cell receptor genes necessary for further lymphocyte development. Consequently, NHEJ deficiency halts lymphocyte development, resulting in B-T-severe combined immunodeficiency in patients and on animial models (4–7). Upon antigen exposure, with T-cell assistance, naïve B cells undergo further gene rearrangement, known as Immunoglobulin heavy chain (IgH) class switch recombination (CSR), to achieve different effector functions. CSR replaces the initially expressed Cµ constant region exons (encoding the IgM antibody) with a set of downstream CH exons encoding different antibody isotypes (e.g., Cγ1 for IgG1). Both NHEJ pathway and the alternative end-joining (Alt-EJ) pathway, which preferentially uses microhomology (MH) at junctions, can mediate CSR. NHEJ deficiencies reduced CSR efficiency by 10% in DNA-PKcs null cells, and upto 50% in cells deficient in core NHEJ factors. The residual junctions mediated by the Alt-EJ pathway (24–30) with MH at the junction and increased chromosomal breaks (31,32). Analyses of V(D)J and CSR efficiency and junctions provide valuable insights on the function of NHEJ factors within a physiological chromosomal context. The core NHEJ factors, including KU, are required for the end-ligation, making them essential for both SJ and CJ formation during V(D)J recombination and CSR (33). As a result, V(D)J recombination is completely abrogated, and CSR is severely compromised in KU-deficient mice (34–38). Conversely, complete loss of DNA-PKcs and Artemis abrogates hairpin opening (and thus CJ formation), but does not abolish end-ligation (including SJ formation and CSR), potentially due to the presence of redundant end-tethering mechanisms (e.g., XLF) (39–42).

While the KU ring formed by the core regions is crucial for KU stability and all known NHEJ functions, KU70 and KU80 have developed additional C-terminal regions in higher eukaryotes (Fig.S1A). The C-terminal domain and tail of KU80 are known to help recruit DNA-PKcs (43,44), but the role of the KU70 C-terminal linker (animo acids, aa 539-556 including the nuclear localization signal, NLS) and SAP domain(aa 560-609) remains unclear (Fig. 1A–B and Fig.S1). In the absence of DNA, the flexible linker allows the SAP domain to take on at least three different positions, including one near the opening of the KU ring (45). Early research indicated that the KU70 SAP domain can bind DNA weakly but consistently (46). However, the SAP domain was not observed in several structural studies including dsDNA(9,10,15,16), leaving its physiological role uncertain. Recent research in *Arabidopsis thaliana*, which lacks a DNA-PKcs equivalent, suggests that the KU70 SAP domain might stabilize KU at DNA ends (47). To elucidate the function of the KU70 SAP domain in mammals, we generated a mouse model with a targeted deletion of amino acids 558-608, encompassing the entire SAP domain while sparing the NLS. The Ku70^ΔSAP^ protein supports the stability of Ku80 and the efficient recruitment of KU to DNA damage sites *in vivo*. Moreover, *Ku70^ΔSAP/ΔSAP^* mice were born at the expected Mendelian ratio, were healthy and exhibited normal lymphocyte development. Both V(D)J recombination and CSR, which entirely depend on KU and NHEJ, remain normal in *Ku70^ΔSAP/ΔSAP^* cells. Structural modeling of KU on long dsDNA suggests that the SAP domain can bind to an adjacent major groove (Fig. 1B), where it might limit KU’s rotation and lateral movement along the dsDNA - a role that might be masked by DNA-PKcs, which caps the end and pushs KU inward. Indeed, in the absence of DNA-PKcs, DNA damage-induced *Ku70^ΔSAP^* foci became significantly weaker. *Ku70^ΔSAP^* compromise the quanlity and quantity of the residual SJs in *DNA-PKcs^−/−^* lymphocytes. At low concentration, Ku70^ΔSAP^ diassociates from the dsDNA ends more readily and at higher concentrations, more Ku70^ΔSAP^ molecules threaded onto the dsDNA. Collectively, these findings revealed a physiological role of the SAP domain in limiting the lateral movement of KU at DNA ends, a function that is otherwise masked by DNA-PKcs.

**Fig. 1.**
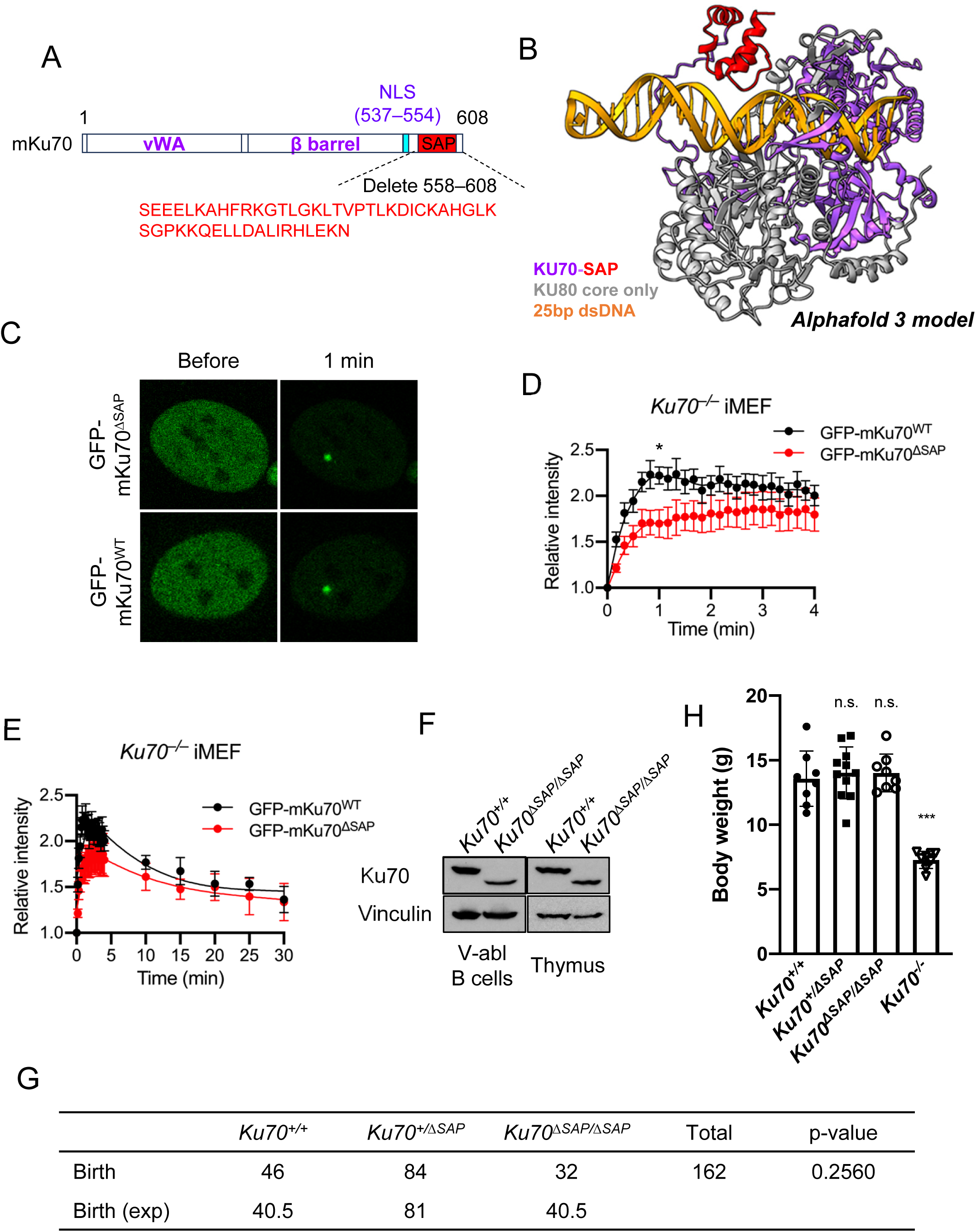
The *Ku70^ΔSAP/ΔSAP^* mice look normal. (*A*) Schematic of mouse Ku70 protein. The nuclear localization signal (NLS) is marked. The SAP domain is in red to be consistent with the structural model in panel *B*. (*B*) The Alphafold 3 generated structural model of full length human KU70, the core region of KU80 (1-545aa) and 25 bp perfect dsDNA complex. The KU70 is colored in purple with red SAP domain. KU80 is in grey and dsDNA is in bright orange. (*C*) Representative images of micro-irradiation-induced GFP-mKu70^WT^ or GFP-mKu70^ΔSAP^ foci in *Ku70^−/−^* iMEFs (34). (*D and E*) The relative intensity of GFP-mKu70^WT^ and GFP-mKu70^ΔSAP^ at DNA damage sites in *Ku70^−/−^* iMEFs. The average ± SEM from at least eight cells were plotted. The star (*: *P* < 0.05) shows the statistics at 1 min post-micro-irradiation. Student’s *t* test was used to calculate the *P* value. This experiment was repeated twice. *D* is for the first 4 minutes and *E* is to 30 minutes. (*F*) Western blot of Ku70ΔSAP in v-abl transformed B cells and thymic cells from indicated mice. (*G*) The number of live born mice obtained at P7 from crosses between *Ku70^+/ΔSAP^* mice. The *P* value was calculated with the chi-squared test. (*H*) The total body weight of adult mice (P21–P24) from the different genotypes indicated. The average ± standard deviation for each genotype were plotted. Student’s *t* test was used to calculate the *P* value. n.s.: no significant difference. ***: *P* < 0.001.

## Results

### Generation of *Ku70^ΔSAP^* allele and mouse models

To investigate the physiological function of the Ku70 SAP domain, we designed a targeting construct to remove part of Exon 13, which encodes the SAP domain (aa 558-608) while preserving NLS and 3’UTR (Fig.S1A-B). The construct also introduced a neo-resistance (NeoR) cassette flanked by FLP recombinase target (FRT) sites into intron 12 (Fig.S1B). The correct targeted allele introduced a new BamHI restriction site, reducing the PacI-BamHI fragment from 12.7 kb in the germline to approximately 5.9 kb. Additional analyses with a NeoR probe confirmed the single integration of the NeoR gene at the designated location, yielding an 8.2 kb fragment (Fig.S1B-C). To determine the impact of the Ku70^ΔSAP^ allele, we generated an N-terminal GFP fused mKu70^ΔSAP^ and control mKu70^WT^ and introduced them into immortalized *Ku70^−/−^* murine embryonic fibroblasts (iMEFs)(34,47,48). Given that *Ku70^−/−^* cells are NHEJ-deficient, we used a micro-irradiation protocol without BrdU sensitization to avoid fluorescence signal saturation (Fig.S1D). Consistent with the SAP domain not being essential for KU stability, mKu70ΔSAP efficiently recruited KU to micro-irradiation sites (Fig. 1C). Although the raise of the relative intensity of DNA damage-induced mKu70^ΔSAP^ foci was delayed, it eventually reached levels comparable to those of mKu70^WT^ (Fig. 1D-E). Additionally, B cell lines and thymus tissue isolated from *Ku70^ΔSAP/ΔSAP^* mice expressed Ku70^ΔSAP^ protein at smaller size but comparable levels (Fig. 1F). In contrast to the severe growth retardation observed in *Ku70^−/−^* mice (49), *Ku70^ΔSAP/ΔSAP^* mice were born at the expected ratio and exhibited normal growth (Fig. 1G-H). Together, these observations suggest that the Ku70 SAP domain is dispensable for murine development. Similar findings were noted on an independently generated Ku70-SAP domain deletion model described in a new preprint (50).

### Normal lymphocyte development and maturation in *Ku70^ΔSAP/ΔSAP^* mice

Given Ku70 is essential for NHEJ and lymphocyte development, we analyzed lymphocyte development and maturation in *Ku70^ΔSAP/ΔSAP^* mice, using the previously characterized *Ku70^−/−^* mice (34,49,51) as control. Flow cytometry analyses show that *Ku70^ΔSAP/ΔSAP^* mice have a normal percentage of naïve (B220^mid^IgM^+^) and recirculating (B220^high^IgM^+^) B lymphocytes in the bone marrow and an expected number of mature T lymphocytes (CD4^+^CD8^−^ and CD4^−^CD8+ single positive, SP) in the thymus (Fig. 2A). In sharp contrast to the complete absence of mature B and T cells in the *Ku70^−/−^* mice, young adult (8–12 weeks) *Ku70^ΔSAP/ΔSAP^* mice have normal counts of splenic IgM^+^ B cells and thymocytes (Fig. 2B-C). The ratio of pre-B cells vs. pro-B cells, a sensitive indicator of successful V(D)J recombination at the IgH loci (52,53), was severely compromised in *Ku70^−/−^* mice but normal in *Ku70^ΔSAP/ΔSAP^* mice (Fig. 2D). Similarly, the frequency of surface TCRβ high thymocytes (Fig. 2A and Fig.S2C), a measurement for successful V(D)J recombination at the TCRα locus, was reduced in *Ku70^−/−^* mice but remained normal in *Ku70^ΔSAP/ΔSAP^* mice. These data suggest that the Ku70 SAP domain is largely dispensable for lymphocyte development and, by extension, V(D)J recombination.

**Fig. 2.**
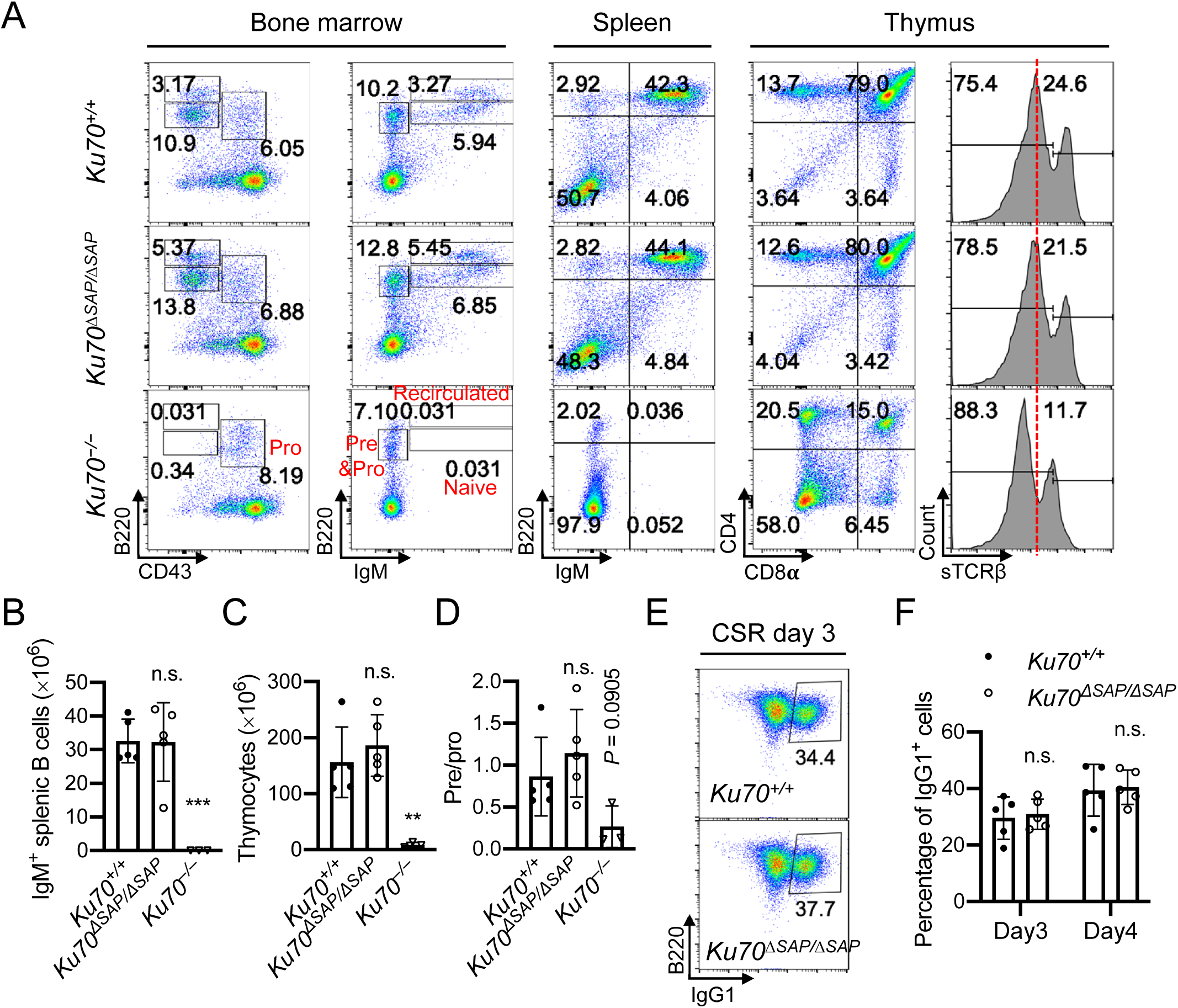
*Ku70^ΔSAP/ΔSAP^* mice have normal lymphocyte development and maturation. (*A*) Representative flow cytometric analyses of bone marrow, spleen and thymus from *Ku70^+/+^*, *Ku70^ΔSAP/ΔSAP^ and Ku70^−/−^* mice. Numbers on the plot are percentages of cells among the gated populations. (*B* and *C*) The total IgM^+^ splenic B cell numbers (*B*) and total thymocyte numbers (*C*). (*D*) The ratio of pre-B (B220^+^IgM^−^CD43^−^) versus pro-B (B220^+^IgM^−^CD43^+^) cells in the bone marrow. (*E*) Representative flow cytometric analyses of IgG1 CSR efficiency after 3 days of cytokine stimulation. The percentage of B220^+^IgG1^+^ cells among all B cells was marked on each panel. (*F*) Quantification of B220^+^IgG1^+^ B-cell percentage on day 3 and 4 of stimulation. For panel *B–D* and *F*, the bars represent the average and the standard deviation of 3 or more mice per genotype. Student’s *t* test was used to calculate the *P* value. n.s.: no significant difference. **: *P* < 0.01, ***: *P <* 0.001.

Next, we tested whether the SAP domain plays a role in IgH CSR. To do this, we isolated and activated *Ku70^+/+^* and *Ku70^ΔSAP/ΔSAP^* CD43^−^ splenic B cells with anti-CD40 and interleukin (IL)-4 to stimulate CSR to IgG1 and IgE. *Ku70^ΔSAP/ΔSAP^* B cells underwent CSR efficiently (Fig. 2E and F), indicating that the Ku70 SAP domain is also dispensable for efficient IgH CSR. *In vivo*, the ratio of antigen experienced recirculated B cells (B220^high^IgM^+^) to naïve B cells (B220^mid^IgM^+^) in the bone marrow was similar in *Ku70^ΔSAP/ΔSAP^* mice and *Ku70^+/+^* ctrl mice (Fig. 2A and Fig.S1F), consistent with normal germinal center responses, including CSR. Furthermore, activating splenic *Ku70^ΔSAP/ΔSAP^* B cells were not sensitive to irradiation (Fig.S1G). Mild end-joining defects might compromise the quanlity of the junction without significantly reducing CSR efficiency, like due to the high GC-content and the repetitive nature of the switch region, which are particularly suitable for Alt-EJ as seen in DNA-PKcs null models (25,30,54). We then analyzed >38,000 unique CSR junctions from three *Ku70^ΔSAP/ΔSAP^* mice along with two *Ku70^+/+^* ctrl mice using high throughput genome-wide translocation sequencing (HTGT-seq)(30,52,54–56) (Fig.S2A-B). We placed the bait primer at the 5’Sμ region and detected genome-wide translocations involving the bait breaks (Fig.S2A). In both genotypes, the vast majority (>90%) of the preys fell in the IgH locus (Fig.S2C). Since most Sε switching are achieved through sequential switching from Sμ to Sγ1, then to Sε, requiring two end-ligations, a preferential reduction of Sε preys has been observed in other NHEJ-deficient backgrounds, including Xrcc4-null, DNA-PKcs null and DNA-PKcs T2609 phosphorylation site mutant mice (30,52,54,57). The IgH preys from both *Ku70^ΔSAP/ΔSAP^* and *Ku70^+/+^* ctrl B cells were evenly mapped into Sμ, Sγ1, and Sε regions, reflecting internal deletion, and CSR to IgG1 and IgE, respectively (58) (Fig.S2D), indicating normal CSR.

A delay in end-ligation kinetics increases the opportunity for end-resection. Since KU has been implicated in end-protection, we also analyzed end-resection, measured by the accumulation of the prey downstream of the core switch regions(30,54,57) and the increased inter-chromosomal and inter-sister CSR with prey at the plus strand (52,58). In both cases, there was no difference between *Ku70^ΔSAP/ΔSAP^* and *Ku70^+/+^* ctrl B cells (Fig.S2 E-H). Moreover, in mutants with mild NHEJ defects, there is often an increase in MH usage at CSR junctions (30,52,54,57), indicating the compensatory use of the Alt-EJ pathway. No changes in MH usage were noted in junctions recovered from *Ku70^ΔSAP/ΔSAP^* B cells (Fig.S2I). Taken together, *Ku70^ΔSAP/ΔSAP^* B cells carried out CSR at normal frequency and with comparable junctional features. We conclude that the SAP domain is dispensable for chromosomal V(D)J recombination and CSR in otherwise wild-type cells.

### Normal lymphocyte development and maturation in *Ku70^ΔSAP/ΔSAP^Xlf ^−/−^* mice

In addition to end-ligation and end-processing, KU has been implicated in end-tethering by providing the anchor points for DNA-PKcs, PAXX, XLF, as well as LIG4(15). Specifically, XLF contributed to two end-tethering pathways: direct anchorage of its C-terminus to KU, and forming a bridge between the two DNA ends via the LIG4-XRCC4-XLF-XRCC4-LIG4 complex (53,59–63). Previous work from we and others showed that loss of XLF revealed a critical role of PAXX and DNA-PKcs in end-tethering, which was otherwise masked by the redundant function of XLF (53,59–63). To investigate whether the Ku70 SAP domain contributes to end-tethering through its DNA binding ability, we generated *Ku70^ΔSAP/ΔSAP^Xlf ^−/−^* mice (28,29). Compared to *Xlf ^−/−^* mice, *Ku70^ΔSAP/ΔSAP^Xlf ^−/−^* mice displayed no additional defects in B and T cell development, as measured by the counts and relative percentage of mature B cells in the bone marrow and spleen, the fraction of DP and SP T cells and the surface TCRβ^high^ T cells in the thymus (Fig.S3A-C). The ratio of Pre/Pro-B and SP/DP T cells are also unaffected (Fig.S3D-E). Furthermore, IgH CSR of *Ku70^ΔSAP/ΔSAP^Xlf ^−/−^* B cells were also comparable to that of *Xlf ^−/−^* ctrl (Fig.S3A and F). These data suggest that the Ku70 SAP domain does not have a redundant end-tethering function that can be unmasked by Xlf loss.

### Loss of DNA-PKcs reveals a role of Ku70 SAP domain in lymphocyte development

The SAP domain exhibits weak but consistent, DNA binding *in vitro* (46). Early X-ray crystal structural analyses of full-length human KU on Y-shaped DNA with ∼14 bp double helix suggest that the SAP domain is positioned from the DNA ends, but failed to reveal its precise location (9). Recent Cry-EM analyses of KU and DNA-PKcs bound to longer dsDNA (30 bp) suggest that the complete footprint of KU extend to ∼20 bp rather than 15 bp (13). Short dsDNA was selected for the structural study because KU is known to slide along DNA, and using short dsDNA facilitates a uniform positioning of KU for structural analysis. To explore whether longer dsDNA would provide a foothold for the SAP domain, we performed structural modeling using AlphaFold 3 with a 25 bp dsDNA. Intriguingly, all five predictions placed the Ku70 SAP domain on the adjacent major groove, away from the DNA ends, with high confidence (Fig. 1B and Fig.S4A-C and Video S1-2). To test the specificity of SAP binding to the major groove, we switched the nucleic acid template from B-form dsDNA to A-form dsRNA, which has a narrower but deeper major groove. Despite similar binding by the ring, on dsRNA, the SAP domain was observed at five different locations, including at the far ends (Fig.S4D). Overlaying the top predictions on dsRNA and dsDNA shows that the KU SAP domain conflicts with the dsRNA helix but fits snugly into the wider major grooves of dsDNA (Fig.S4E). Given that A-form dsRNA helix is wider and shorter than B-form dsDNA with the same sequence, we also tested a 39 bp longer dsRNA, which similarly left the SAP domain near the ends (Fig.S4F). Together, these modeling results support a length-independent, selective binding of the KU SAP domain to the major groove of DNA (Fig. 1B and Fig. S4A). By binding to the adjacent major groove, the Ku70 SAP domain could potentially restrict KU’s rotation and limit the lateral movement of the KU ring on DNA, as first speculated by Goldberg and Walker in the discussion (9). If so, the recruitment of DNA-PKcs, which interacts with Ku70 and caps the ends, might mask this role of the SAP domain in limiting KU’s lateral movement on DNA (10,13). Consistent with this hypothesis, in *Arabidopsis thaliana,* which lacks DNA-PKcs, loss of SAP domain reduces KU loading on DNA (64), presumably by allowing the KU ring to slide off the DNA ends more easily. If this model is correct, the absence of DNA-PKcs would reveal a role for the Ku70 SAP domain in NHEJ.

To test this hypothesis, we generated *Ku70^ΔSAP/ΔSAP^DNA-PKcs^−/−^* mice (42). DNA-PKcs is essential for the recruitment and activation of Artemis endonuclease for end-processing, including CE hairpin opening crucial for V(D)J recombination(14,42,65). Consequentially, CJ formations and lymphocyte development are largely abrogated in *DNA-PKcs^−/−^* mice, while the joining of the blunt SJ is only moderately affected (41,42,66). However, a small fraction of *DNA-PKcs^−/−^* thymocytes can leak through, resulting in an apparent TCRβ^high^ T cell population in the thymus (41,42) (Fig. 3A). Notably, *Ku70^ΔSAP/ΔSAP^DNA-PKcs^−/−^* mice were born at the expected Mendilian ratio and were statistically significantly smaller than the age-matched *DNA-PKcs^−/−^* ctrls, suggesting additional end-joining defects (Fig.S5A-B). The introduction of *Ku70^ΔSAP/ΔSAP^* also futher reduced cellularity (Fig. 3B) and eliminated leaky thymocyte development as shown by the absence of TCRβ^high^ T cells in the thymus, and SP T cells in the spleen (Fig. 3A and 3C-D and Fig.S5C). Although not apparent from flow cytometry, the absolute numbers of IgM+ B cells in the bone marrow and spleen were also decreased in *Ku70^ΔSAP/ΔSAP^DNA-PKcs^−/−^* mice compared to *DNA-PKcs^−/−^* ctrl mice (Fig. 3B, and 3E-F). These findings are consistent with a role for the Ku70 SAP domain in lymphocyte development that is otherwise masked by DNA-PKcs.

**Fig.3.**
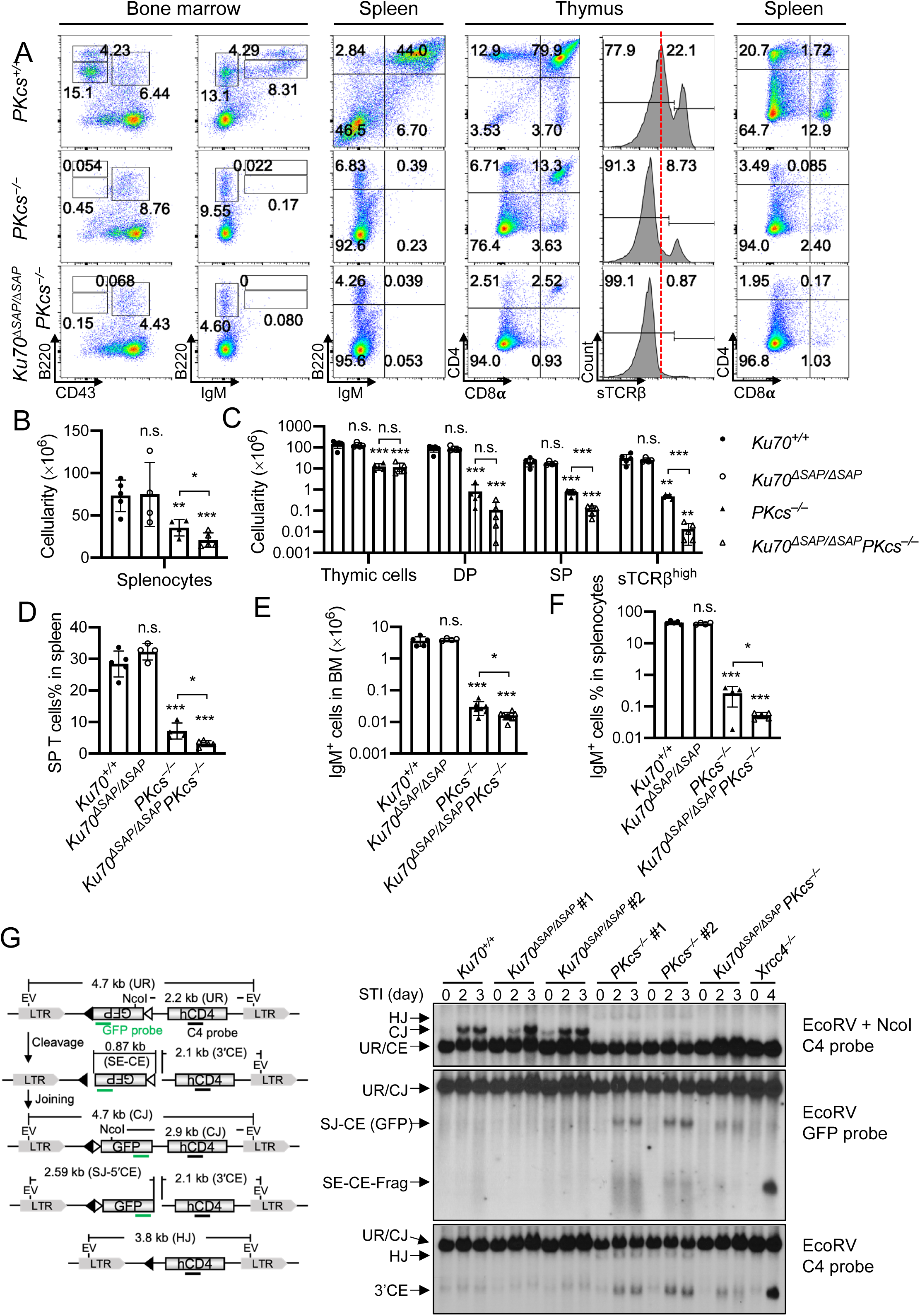
SAP domain of Ku70 contributes to lymphocyte development in *DNA-PKcs^−/−^* mice. (*A*) Representative flow cytometric analyses of bone marrow, spleen and thymus from *DNA-PKcs^+/+^*, *DNA-PKcs^−/−^ and Ku70^ΔSAP/ΔSAP^DNA-PKcs^−/−^* mice. Numbers on the plot are percentages of cells among the gated populations. (*B*) The total splenic cell numbers. (*C*) The total thymocyte numbers, CD4^+^CD8^+^ DP immature T-cell numbers, the sum number of CD4^+^ or CD8^+^ SP T-cells and surface-TCRβ^+^ (the right-side peak in panel *A*) cell numbers in thymus. (*D–F*) The percentage of SP T-cells in spleen (*D*), IgM^+^ cells in bone marrow (BM) (*E*) and IgM^+^ cells in spleen (*F*). For panels *B–F*, the bars represent the average and the standard deviation of 4 or more mice per genotype. Student’s *t* test was used to calculate the *P* value. n.s.: no significant difference. *: *P* < 0.05, **: *P* < 0.01, ***: *P <* 0.001. (*G*) Schematic of pMX-INV chromosomal V(D)J recombination substrates. The pMX-INV vector has a major pair of recombination signal sequences (RSSs) flanking the inverted GFP cassette. LTR=long terminal repeat from the retrovirus vector, hCD4=truncated hCD4 cDNA encoding the transmembrane and extracellular domains of human CD4, IRES=Internal Ribosome Entry Sites, filled triangle= 5′ 12-spacer RSS, open triangle = 3′ 23-spacer RSS, and EV=EcoRV cutting site. The diagram includes unrearranged substrate (UR), signal end (SE) intermediates, coding end (CE) intermediates, 4.7 kb coding joint (CJ), 2.59 kb signal joint (SJ) and 3.8 kb hybrid joint (HJ) products. Canonical V(D)J recombination between 12 and 23 RSSs inverts the GFP cassette and forms an SJ and a CJ. On the right, the Southern blot results of EcoRV-NcoI-double-digested (top) or EcoRV-digested (bottom) DNA with the C4 probe (top) or GFP probe (bottom). The genotypes and the days of STI treatment were marked on the top. The identities of the products were marked on the left.

### Loss of the Ku70-SAP domain alters the V(D)J recombination junctions isolated from *DNA-PKcs^−/−^* B cells

To determine whether the KU70 SAP domain contributes to lymphocyte development at the step of chromosomal V(D)J recombination, we derived v-abl kinase transformed immature B cells (v-abl cells) from control wild-type, *Ku70^ΔSAP/ΔSAP^* , *DNA-PKcs^−/−^*, and *Ku70^ΔSAP/ΔSAP^DNA-PKcs^−/−^* mice, all of which carried the EµBcl-2 transgenes (17,53,67,68). Treatment of these v-abl cells with STI571/Gleevec, a v-abl kinase inhibitor, arrests cells in G1 and induces RAG recombinase expression, resulting in efficient chromosomal V(D)J recombination of integrated V(D)J recombination substrates (*e.g.*, pMX-INV) (68). The Bcl-2 transgene is necessary to obviate the apoptosis induced by STI571(68). Multiple v-abl cell lines were generated from each genotype, each harbored a chromosomal integrated inverted V(D)J recombination substrate (pMX-INV) designed to assay both CJs and SJs (Fig. 3G). Given the essential role of DNA-PKcs in hairpin opening and CJ formation, we focused on the impact of KU70 SAP domain on SJs and SEs end-ligation. Successful V(D)J recombination inverts the GFP ORF and forms a 5’SJ and a 3’CJ (68) (Fig.3G). As such, the GFP will be in the same transcriptional orientation as the LTR promoter, leading to GFP expression. Due to hairpin opening defects, *DNA-PKcs^−/−^* and *Ku70^ΔSAP/ΔSAP^DNA-PKcs^−/−^* lines cannot generate substantial CJs after STI treatment and accumulated high levels of free CEs and the SJ-CE fragments containing the GFP ORF (Fig. 3G). Consistent with the developing data, the result suggests that the loss of Ku70 SAP domain moderately, but surely, attenuates SJ-CE accumulation in *DNA-PKcs^−/−^* cells, measured both decrease intensity and degradation of the SJ-CE products. But the level of SEs seems insufficient to explain the much more severe developmental defects in *Ku70^ΔSAP/ΔSAP^DNA-PKcs^−/−^* mice vs. the *DNA-PKcs^−/−^* controls (Fig. 3A-E).

In addition to joining efficiency, reduced fidelity could also compromise lymphocyte development *in vivo*. To determine whether the quality of the SJs formed in *Ku70^ΔSAP/ΔSAP^DNA-PKcs^−/−^* cells has changed, we performed HTGT-seq (38,55) using a bait primer at the 5’SE (Fig. 4A). The results revealed thousands of individual joins between a 5’SE and a various genome-wide joining partners (preys), including the bona fide partner 3’SE (Fig. 4A and Fig.S5D). In addition, HTGT-seq can reveal the orientation of the joining product. We defined junctions in the 5’ to 3’ direction as “plus” (in blue) and those in the 3’ to 5’ direction relative to the substrate as “minus” (in red). In both WT and *Ku70^ΔSAP/ΔSAP^* lines, the vast majority (∼80%) of the preys for the 5’SE bait are the canonical 3’SEs (in minus orientation – red) within the pMX-INV substrate, indicative of faithful SJs (Fig. 4*A* and Fig.S5E-F). *DNA-PKcs^−/−^* cells exhibited a significantly increased fraction of translocations outside the pMX-INV, consistent with a crucial role of DNA-PKcs in end-tethering and end-ligation (Fig.S5E). The hybrid joins (HJ) formed between 5’SEs and 3’CEs, indicated by prey in the plus (blue) orientations, were rare in WT and *Ku70^ΔSAP/ΔSAP^* lines (< 2%) but common (33.83%) in *DNA-PKcs^−/−^* cells consistent with delayed hairpin opening and the blend of CEs with SEs (Fig. 4*A*). Notably, loss of Ku70 SAP domain significantly reduced HJ formation (from 33.83 to 6.03%) in *DNA-PKcs^−/−^* cells (Fig. 4*A*), indicating a role of the SAP domain in end-ligation in *DNA-PKcs^−/−^* cells. We further analyzed the fidelity of SJs recovered from WT, *Ku70^ΔSAP/ΔSAP^*, *DNA-PKcs^−/−^*, and *Ku70^ΔSAP/ΔSAP^DNA-PKcs^−/−^* lines. While the vast majority of the SJs were precise in WT and *Ku70^ΔSAP/ΔSAP^* lines, SJs from *DNA-PKcs^−/−^* lines exhibited frequent and larger deletions, with a mean size of ∼15 nt, coinciding with the footprint of KU (Fig. 4B). Interestingly, the loss of the SAP domain decreased the deletion size from 15 nt back to 3 nt (Fig. 4B). Base pair resolution analyses of bait-side deletions from bona fide SJs showed that the loss of the SAP domain alone (*Ku70^ΔSAP/ΔSAP^*) has no impact on deletion (similar to WT) (Fig. 4C-D). *DNA-PKcs^−/−^* cells frequently exhibited deletions, especially large deletions (> 20 bps), which were significantly reduced in *Ku70^ΔSAP/ΔSAP^* background (Fig. 4C-D). This data suggests that the loss of the KU70 SAP domain reduces overall ligation efficiency while allows the join to occur closer to the DNA ends in *DNA-PKcs^−/−^* cells. A similar decrease of deletion was noted even when all junctions (regardless of prey) were pooled together (Fig.S5D). Given the KU ring occupies about 15-20 bp of DNA, the data is consistent with a model in which, in the *DNA-PKcs^−/−^* cells, joining typically occurs after endonuclease-mediated resection that removes KU, whereas in *Ku70^ΔSAP/ΔSAP^DNA-PKcs^−/−^* cells, *Ku70^ΔSAP^* slides off more easily, allowing ligation to occur closer to the DNA ends.

**Fig. 4.**
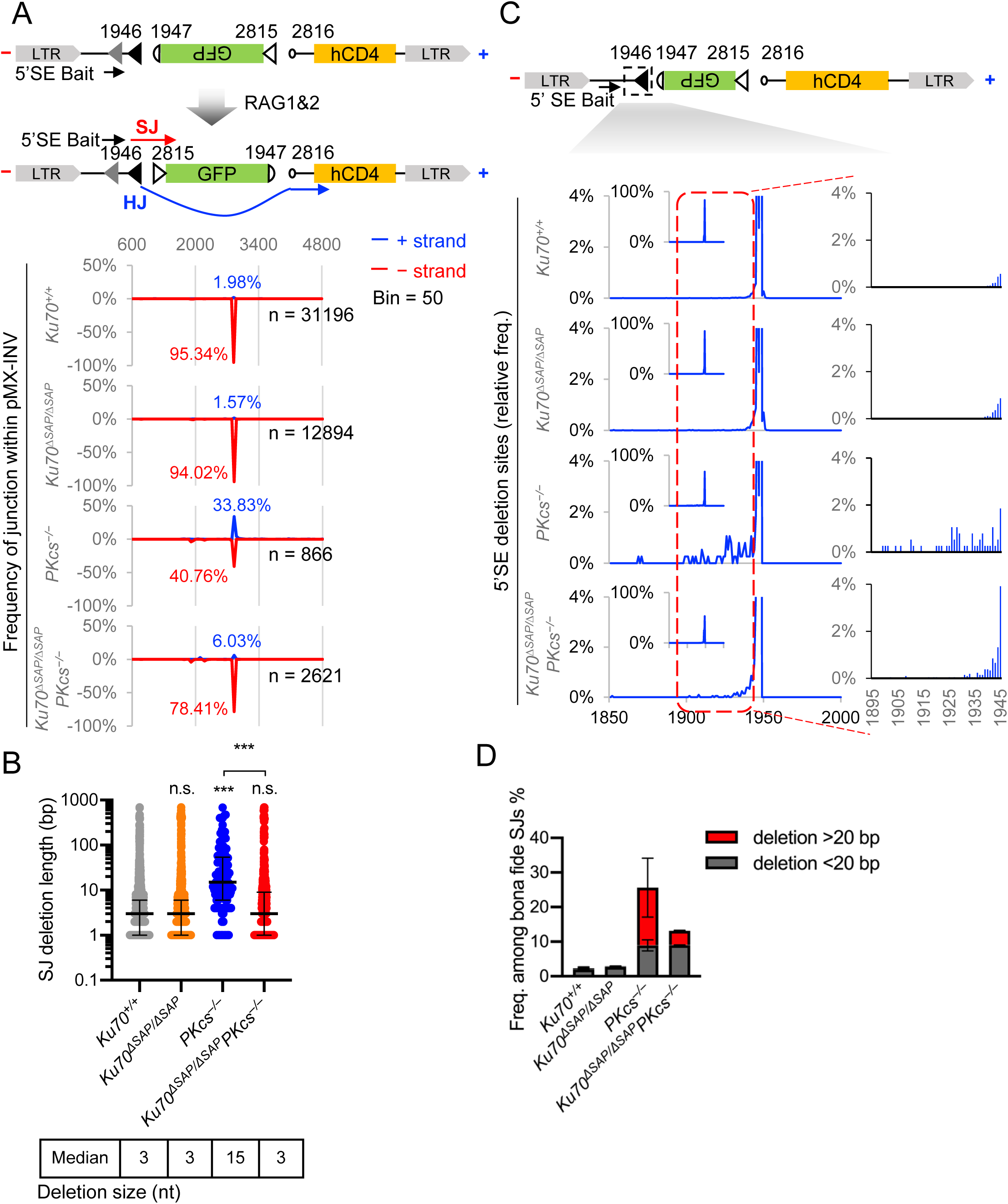
Ku70-SAP domain alters the V(D)J recombination junctions isolated from *DNA-PKcs^−/−^* B cells measured by HTGT-seq analyses using a SE bait break on pMX-INV substrate. (*A*) (Top) The diagram of the possible junctions recovered using a 5′SE bait. The location of canonical 5′ 12 RSS (black filled triangle), alternative pseudo 5′ 12RSS (grey triangle) and 3′ 23 RSS (open triangle), 5′CE (semicircle near 1947), and 3′CE (circle near 2816) are marked. (Bottom) The spatial distribution of all preys (50 bp bin) is plotted against the location on pMX-INV substrate. The red traces indicate preys aligned to the (−) strand (canonical SJ products) and the blue traces indicate preys aligned to the (+) strand (HJ products). The total number of junctions is shown on the right. (*B*) The distribution of total SJ deletion length (bp). Deletions from both sides of the breaks were added together. The lines represent median and interquartile range. Mann-Whitney test was used to calculate the *P* value. n.s.: no significant difference. ***: *P <* 0.001. The pool of all data from each genotype (see *Fig.*S5D for sample information) was plotted. (*C*) (Top) Schematic diagram of pMX-INV substrate for V(D)J recombination with the location of the 5′ bait SE RSSs (the filled black triangle). (Bottom) The spatial distribution of 5′SE terminal locations among all bona fide SJs within pMX-INV substrate (1 bp bin). (*D*) The frequency of 5′ bait SE with deletion larger or smaller than 20 bp among all bona fide signal junctions recovered in each genotype. The bars represent the average ± standard deviation of two samples per genotype.

### Loss of the Ku70-SAP domain does not alter the CSR junctions isolated from *DNA-PKcs^−/−^* B cells

Next, we analyzed IgH CSR. To circumvent the requirement for DNA-PKcs in B cell development, we introduced pre-assembled IgH and IgL genes into the germline of the mice (HL^K^)(69,70). Using this approach, we and others have characterized the role of DNA-PKcs, Lig4 and other essential NHEJ factors in CSR (25,30,37,71). To analyze CSR, we isolated and activated *HL^K/K^*, *DNA-PKcs^−/−^HL^K/K^*, and *Ku70^ΔSAP/ΔSAP^DNA-PKcs^−/−^HL^K/K^* B cells with anti-CD40 and IL-4 to stimulate CSR to IgG1 and IgE. As expected, *DNA-PKcs^−/−^HL^K/K^* B cells underwent CSR at ∼80% efficiency of the *HL^K/K^* ctrls and *Ku70^ΔSAP/ΔSAP^DNA-PKcs^−/−^HL^K/K^* B cells shown no further reduction in CSR efficiency (Fig. 5A-B). Next, we analyzed >5,000 unique switch junctions from at least two mice of each genotype (Fig.S6A). We placed the bait primer at the region and detected genome-wide translocations involves 5’Sμ (Fig. 5C-D). In all genotypes, the vast majority (>90%) of the preys were mapped within IgH (Fig.S6B). Within IgH, the preys were evenly distributed into Sμ, Sγ1, and Sε in controls, but there were significantly fewer Sε preys in *DNA-PKcs^−/−^HL^K/K^* and *Ku70^ΔSAP/ΔSAP^DNA-PKcs^−/−^HL^K/K^* B cells, consistent with end-joining defects (30,54,57) (Fig.S6C). Two other indications of end-ligation defects are increasing end-resection, measured by increased preys outside the core switch regions (30,54,57) and the increased inter-chromosomal and inter-sister CSR with preys on the plus strand (58). For Sμ, Sγ1 and Sε preys, both *DNA-PKcs^−/−^HL^K/K^* and *Ku70^ΔSAP/ΔSAP^DNA-PKcs^−/−^HL^K/K^* B cells displayed similar increased resection compared to the *HL^K/K^* control (Fig. 5 C-E and Fig.S6D-E). There was a significant shift to MH usage at CSR junctions in both *DNA-PKcs^−/−^HL^K/K^* and *Ku70^ΔSAP/ΔSAP^DNA-PKcs^−/−^HL^K/K^* B cells, but no difference in between the two (Fig. 5F). We conclude that, given the high GC content of the substrates and the scattered AID-initiated breaks, the Ku70 SAP domain has no measurable impact on the quality or quantity of CSR junctions. This finding is consistent with the small footprint of KU (∼15-20 bp) that can be masked by the multiple breaks generated by AID in the repetitive switch regions.

**Fig. 5.**
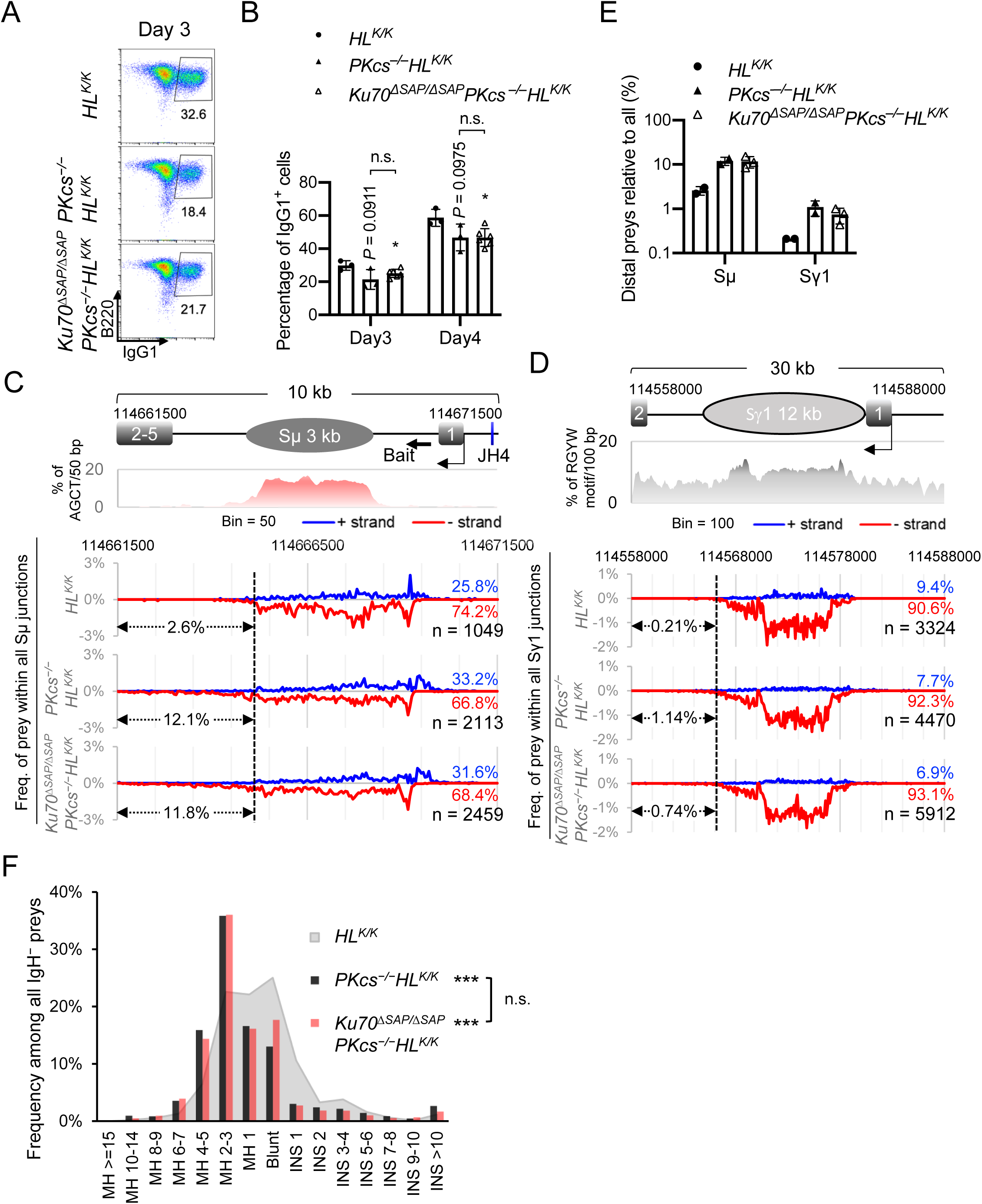
Ku70-SAP domain does not alter the CSR junctions isolated from *DNA-PKcs^−/−^* B cells. (*A*) Representative flow cytometric analyses of IgG1 CSR efficiency after 3 days of cytokine stimulation. The percentage of B220^+^IgG1^+^ cells among all B-cells was marked on each panel. (*B*) Quantification of B220^+^IgG1^+^ B-cell percentage on day 3 and 4 of stimulation. The bars represent the average and the standard deviation of 3 or more mice per genotype. Student’s *t* test was used to calculate the *P* value. n.s.: no significant difference. *: *P* < 0.05. (*C*) The spatial distribution of preys in Sμ region. Schematic of the Sμ region with the bait site at the 5′Sμ marked by an arrow is at Top. The frequency of AGCT motifs per 50-bp double-strand DNA is plotted. The values between y-axis and the dashed line (114,661,500∼114,665,100) indicate the percentage of preys that fall outside the core Sμ region. (*D*) The spatial distribution of preys in Sγ1 region. Schematic of the Sγ1 region is at Top. The frequency of RGYW motifs per 100-bp double-strand DNA is plotted. The values between y-axis and the dashed line (114,558,000∼114,566,000) indicate the percentage of preys that fall outside the core Sγ1 region. For panels *C* and *D*, the pool of all data from each genotype was plotted (see *Fig.*S6*A* for sample information). All (−, red, below zero) and (+, blue, above zero) strand preys add up to 100%. For each genotype, the percentage of (+) strand (blue) and (−) strand (red) junctions and total number are marked at the right. (*E*) Relative frequency of distal preys among the preys in Sμ and Sγ1 region. (*F*) The distribution of all IgH junctions by junction type (MH, microhomology. INS, insertion). This graph represents the pool of at least two independent libraries (see *Fig.*S6A for sample information) of each genotype. Kolmogorov-Smirnov test was used to calculate the *P* value. n.s.: no significant difference. ***: *P* < 0.001.

### Ku70-SAP domain is critical for KU-DNA binding *in vitro*

If the Ku70-SAP domain limits lateral movement of KU on DNA ends in the absence of DNA-PKcs, the model would predict that Ku70^ΔSAP^ falls off the DNA easily at low concentrations and accumulates on the DNA at high concentrations. To test this, we purified KU^ΔSAP^ and KU^WT^ and measured its fficient with Surface Plasmon Resonance (SPR). The result revealed that KU^ΔSAP^ has a moderate and consistent reduced affinity to dsDNA (Fig. 6A-B). Moreover, KU70^ΔSAP^ disassociated from the DNA more readily then KU^WT^ (Fig. 6C) and could be easily competed off by KU^WT^ (Fig. 6D and Fig.S7A-C). While the above data supports a role for the SAP domain in promoting DNA binding, if the SAP domain indeed limits the lateral movement of KU, the model would also predict that at high concentrations, more KU ring containing the KU70^ΔSAP^ might be loaded onto the dsDNA. Thus, we performed a gel-shief assay with a range of KU concentration. Indeed, KU^ΔSAP^ is less efficient at occupying both ends of the dsDNA at low concentration while at high concentration (*e.g.*, 35 nM), it is easier for more KU^ΔSAP^ to be loaded on to the dsDNA, consistent with increasing freedom in lateral movement (Fig. 6E). How about *in vivo*? We tested the formation of DNA damage induced Ku70^ΔSAP^ and Ku70^WT^ foci in *DNA-PKcs^−/−^* iMEFs. In contrast to the similar recruitment and kinetics observed in *DNA-PKcs^+/+^* iMEFs (Fig. 1D-E), in *DNA-PKcs^-/-^* iMEFs, the GFP-Ku70^ΔSAP^ foci were significantly weaker than those of GFP-Ku70^WT^ (Fig. 6F-G and Fig.S7C), consistent with the observation in *Arabidopsis thaliana* without a DNA-PKcs homolog (64). Together, this data supports a model in which the SAP domain of Ku70 binds to DNA and limits the lateral movement of the KU ring on the DNA. In mammals, this function of the Ku70-SAP domain can be masked by the presence of DNA-PKcs, which caps the ends (Fig. 6H). Our study provides the first physiological function of KU70-SAP domain *in vivo*.

**Fig. 6.**
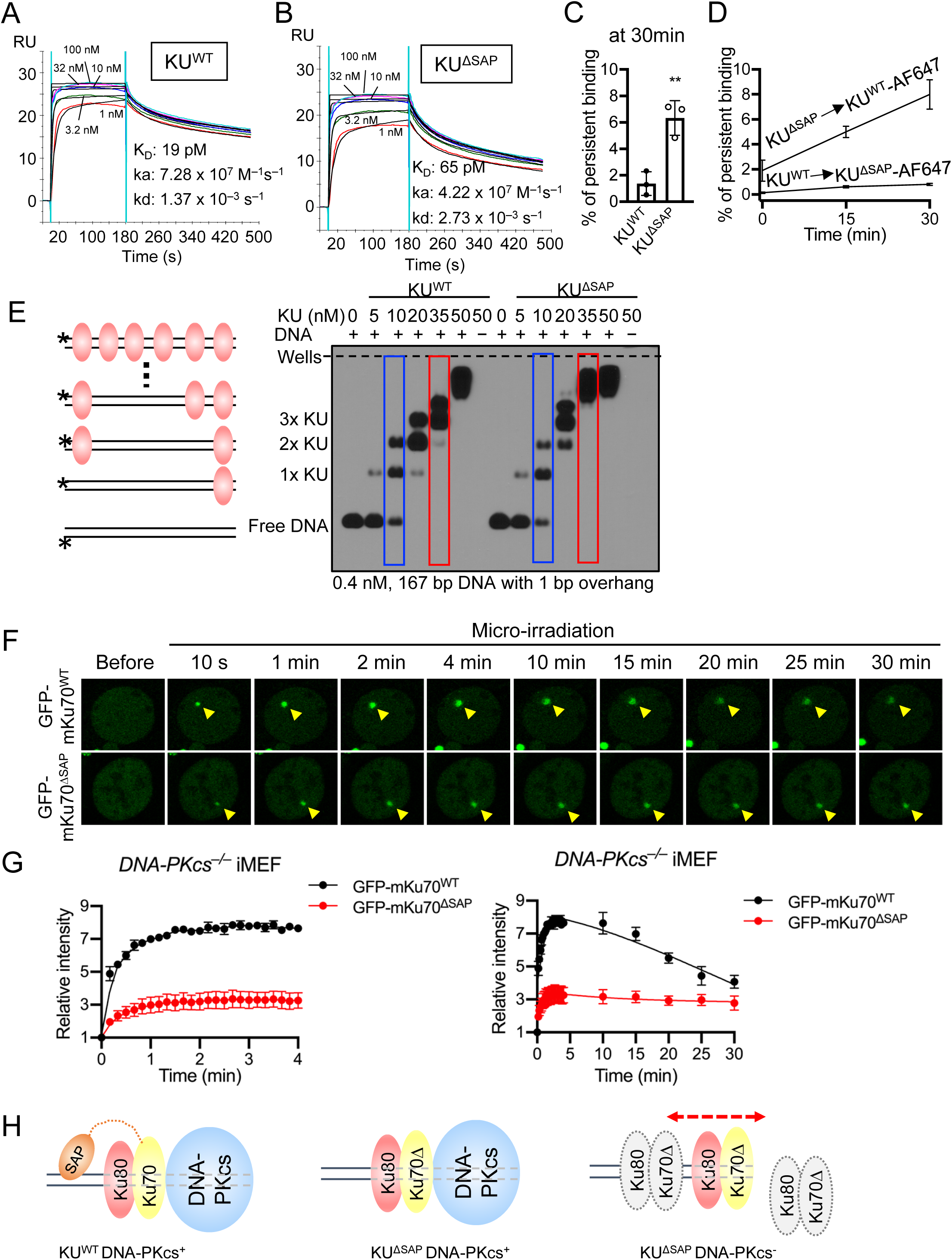
Ku70-SAP domain is critical for KU loading *in vitro*. (*A* and *B*) Indicated concentrations of KU^WT^ (*A*) or KU^ΔSAP^ mutant (*B*) were applied to a sensor chip coated with 400 bp DNA (blunt end) on Biacore T200. Curves (black) proximate to each colored curve indicating different concentrations of KU are fitted curves as a mathematical model generated on Biacore’s software. (*C*) Bar graph of percentage of persistent binding of AF647 labeled KU WT and ΔSAP. The error bar represents standard deviation from three replicates. The percentage of persistent binding between KU^WT^ and KU^ΔSAP^ in the exchange assay is significantly different (*P* value = 0.00562). (*D*) Time course plot of displacement of (top) KU^ΔSAP^ by AF647 labeled KU^WT^ and displacement of (bottom) KU^WT^ by AF647 labeled KU^ΔSAP^. Error bars represent standard deviation from three replicates. (*E*) Indicated amounts of KU WT or ΔSAP mutant were incubated with 167 bp DNA with 1 bp overhang, and analyzed on membrane transferred from agarose gel. Two KU can bind to the dsDNA (at each end) without the need for double loading or sliding. (*F*) Representative images of micro-irradiation-induced GFP-mKu70^WT^ or GFP-mKu70^ΔSAP^ foci in DNA-PKcs knockout iMEF cells. The yellow arrowheads point to the area of micro-irradiation. (*G*) The relative intensity kinetics of GFP-mKu70^WT^ and GFP-mKu70^ΔSAP^ at DNA damage sites in DNA-PKcs knockout iMEF cells within 4 minutes (left) and 30 minutes (right). The average ± SEM from at least eight cells were plotted. This experiment was repeated twice. (*H*) the diagram shows that DNA-PKcs can cap the ends to prevent Ku lateral movement. In the absence of DNA-PKcs and SAP domain, KU can rotate on DNA and produce a lateral movement-fall off the DNA at low concentration and load multimers at high concentration.

## Discussion

KU binding represents a commitment to the NHEJ pathway. While the ring formed by the core domains of KU70 and KU80 is responsible for end-sensing and critical for end-protection, end-tethering, end-processing, and eventually end-ligation, the role of the SAP domain of Ku70 remains elusive. Due to the flexible linker, structure analyses revealed multiple possible configurations of the SAP domain in Apo and DNA-bound forms, ranging from the aperture to the side (45). Using genetically engineered mouse models, our study thoroughly investigated the role of the Ku70 SAP domain in mammalian NHEJ using lymphocyte development as the model system. Our results showed that in otherwise wild-type cells, the Ku70 SAP domain is dispensable for the recruitment of KU and the efficient repair of DSBs generated during V(D)J recombination and CSR. Moreover, using high throughput analyses of DNA repair junctions, we further showed that the Ku70 SAP domain is also dispensable for the precise joining of blunt signal ends. Furthermore, using Xlf-deficient mice with compromised end-tethering as the background, we excluded a role of Ku70 SAP domain in end-tethering and NHEJ even in sensitive backgrounds. Careful analyses of the early and recent structure of KU on DNA prompted us to perform a structural modeling with AlphaFold 3 with a long dsDNA substrate. The results consistently placed the SAP domain on the adjacent major groove on dsDNA (but not dsRNA). Based on this model, we hypothesized that the SAP domain might restrict the rotation of the KU ring on DNA and prevent the lateral movement of KU. Indeed, both genetic and biochemical evidence support this model. Specifically, removing DNA-PKcs that caps the DNA ends and prevents lateral movement of KU revealed a critical role of the SAP domain in promoting end-ligation and the stable association of KU at the DNA ends. Moreover, an independent study found that in A*rabidopsis thaliana,* which does not have a DNA-PKcs homolog, the Ku70 SAP domain stabilizes KU at the DNA ends (64). Biochemically, at lower KU concentrations, the SAP domain promotes KU binding to DNA, presumably by preventing KU from sliding off. At high KU concentrations, the SAP domain limits the number of KU rings that can be threaded onto the DNA, again by limiting lateral movement of KU. Moreover, in DNA-PKcs deficient cells, KU protects the ends from exonuclease resection and helicase activity. As such, residual junctions often contain a deletion of >15-20 bp consistent with endonuclease resection to remove the KU ring and allow end-ligation via a micro-homology mediated pathway. In this context, the loss of the SAP domain destabilizes the DNA-damage-induced KU foci measured by quantitative live cell imaging and allows direct end-ligation without the need for extensive end-resection (presumably to remove KU). Indeed, the base pair deletion in SJs recovered from *Ku70^ΔSAP/ΔSAP^DNA-PKcs^−/−^* cells is much smaller than those found in *DNA-PKcs^−/−^* cells. Consistent with this model, loss of KU rescued the embryonic lethality of Lig4-deficient mice (72) and reduced deletion sizes and microhomology usage in CSR junctions recovered from *Lig4^−/−^* B cells (37). Taken together, our study identified a physiological function of the Ku70 SAP domain in limiting the lateral movement of KU on dsDNA that is partially masked by DNA-PKcs (Fig. 6H). Consistent with a role in limiting, but not eliminating lateral movement of KU which is necessary for DNA-PKcs loading and end-processing, the DNA-binding capability of SAP has been demonstrated by southwestern blotting (46), but is too weak to detect in a gel-shift assay and seems specific to the major grooves of dsDNA (rather than dsRNA).

When did Ku70 acquire the SAP domain? The yeast Ku70 does not have a well-folded SAP domain (Fig. S1), instead, it just has a flexible linker with a likely NLS signal. In plants (e.g., A*rabidopsis thaliana*) and insects (e.g., *Drosophilae*), Ku70 acquires the SAP domain (64) before the emergence of DNA-PKcs. In there, the SAP domain likely plays an important role in stabilizing KU on the DNA ends and promotes NHEJ. How about in vertebrates and mammalian cells? While the presence of and the recruitment of DNA-PKcs can cap the DNA ends and limit the inward rotation of KU by directly interacting with Ku70, several evidences suggest that DNA-PKcs has to be released before end-ligation (10,14,17). The release of DNA-PKcs requires its own kinase activity and potentially the accumulative phosphorylation of DNA-PKcs by itself and the related ATM kinase *in vivo*. Accordingly, we showed that the presence of kinase dead DNA-PKcs completely blocks end-ligation in a KU-dependent manner (17,30). Single-molecule studies and Cryo-EM studies identified a long-range NHEJ complex with DNA-PKcs that is not compatible with end-ligation and a short-range ligation complex without DNA-PKcs (15,18). Based on those findings, we speculate that Ku70 SAP domain might have a conserved role in stabilizing KU on the DNA ends in the transient short-range NHEJ complex during end-ligation. Notably, in contrast to species without DNA-PKcs, the loading of DNA-PKcs pushes the KU ring inward ∼10 bp away from the DNA ends (10,13), which might explain the lack of devastating NHEJ defects in the *Ku70^ΔSAP/ΔSAP^* mice with DNA-PKcs compared to the corresponding A*rabidopsis thaliana* without DNA-PKcs homolog (64). The multi-faceted interaction of KU with Xlf, Paxx, and Lig4/Xrcc4 in the short-range complex might further limit KU rotation and prevent KU from sliding off. Accordingly, we did not detect cellular radiation sensitivity (Fig. S1G) or V(D)J recombination defects in *Ku70^ΔSAP/ΔSAP^* B cells in our study (Fig.2 and 4). A preprint reports that a comparable Ku70-SAP deficient mouse model also has no V(D)J recombination or lymphocyte development defects despite a moderate whole body radiation sensitivity (50).

If the SAP domain binds to the DNA consistently, why has the SAP domain not been seen in previous structural studies? This might have to do with the length of DNA substrates provided. As mentioned, the early X-ray crystal structural analyses used a Y-shaped DNA with only 14 bp dsDNA. The choice was driven by the need to isolate a consistent structure, matching the substrate to the estimated minimal footprint of KU at the time (∼14-15bp dsDNA) to prevent multiple KU from loading on longer DNA (9). Yet, with this short dsDNA substrate, an additional major groove is not available for the SAP domain to bind (Fig.1B). When longer dsDNA (30 bp) was provided to study the loading of KU and DNA-PKcs, DNA-PKcs pushed KU inward and occupied the additional 10 bp and, as such, additional major grooves were still not available (10,13,15,16). This choice might be necessary for structural analyses, given the ability of KU to slide on long dsDNA substrates. But using AlphaFold 3, we showed that a longer dsDNA (>25 bp) and the additional dsDNA major grooves are required to place the Ku70 SAP domain on the adjacent major groove. The actual configuration of Ku70 SAP will have to await future validation via structural analyses.

Taken together, we provided both genetic and biochemical data as evidence to support a role of the Ku70 SAP domain in limiting lateral movement of KU that is partially masked by DNA-PKcs in mammalian cells. These results highlight the redundancy and dynamic assembly of the mammalian NHEJ complex and its role in regulating timely and precise DSB repair during lymphocyte development as well as in somatic tissues.

## Acknowledgment

We thank the members of the Zha Lab for helpful discussions and advice on lymphocyte development and sequence analyses. Due to space limitations, we could not cite all the original publications and cite reviews when necessary. This work was supported by NIH R01CA275184 to SZ and ER; 5R01CA158073 and P01CA174653 to SZ; ANR AAPG2023 - PRC – XXL to MM. YZ is a fellow of Cancer Research Institute. This research was funded in part through the NIH/NCI Cancer Center Support Grant P30CA013696 to the Herbert Irving Comprehensive Cancer Center (HICCC) of Columbia University.

## Author Contributions

Y.Z. and S.Z. designed the research; Y.Z. generated and characterized the Ku70delSAP animal models and collected and analyzed the HTGT-seq data. B.J.L. performed Southern analyses of the v-abl kinase transformed cells, S.F and M.M purified the KU70,= KU70delSAP, and DNA—PKcs protein and performed the SRP and EMSA. S.J. performed the binding and FRET analyses in E.R.’s lab. H.Z and A.L. contributed to the cell biology analyses. K.W. performed Alphafold 3 modeling and generated the related figures. Y.Z. and S.Z. analyzed data. S.Z. and Y.Z. wrote the paper with help from other authors.

## Materials and Methods

### Mice

*Ku70^+/−^*(48), *DNA-PKcs^+/−^*(*42*), *Xlf ^+/−^* (29,73), *HL^K^*(69,70) and *Eμ-Bcl2^+^* (74) mice have been described previously. The Ku70^ΔSAP^ allele deletes the C-terminal SAP domain (558∼608 amino acids, corresponding to 560∼609 in human KU70) using homology targeting (Fig. 1A, SI Appendix. Fig. S1B). Briefly, the pEMC-based targeting construct was used to insert a neomycin-resistance (NeoR) cassette in intron 12, about 400 bp upstream from the truncation site in exon 13 of the murine Ku70 (*Xrcc6*) locus. The 5′ arm is ∼3.5 kb and the 3′ arm is ∼2.9 kb. We deleted the SAP domain (153 bp) in the 3′ arm by overlapping PCR (SI Appendix. Fig. S1B). CSL3 ES cells (129/sv background) were targeted by electroporation and screened via PCR using primers flanking the 3′ arm (5′-GCCTGAAGAACGAGATCAGC-3′ and 5′-CCACCTCACCTGGCTTTTTA-3′). The correct clones were confirmed by Southern blotting (PacI and BamHI digestion, with the 5′ probe generated by PCR using primers 5′-CCAGCTTGAATGTATGAAACCCT-3′ and 5′-CCTTGTGTCCTCTAAGCCT-3′, and Neo probe) (SI Appendix. Fig. S1B-C). The expected germline (GL) band is ∼12.7 kb, and the targeted band is ∼5.9 kb (SI Appendix. Fig. S1C). Six targeted clones were sequenced and verified to have the desired Ku70^ΔSAP^ truncation. One of them was injected for germline transmission. The chimeras were bred into the Rosa26FLIP/FLIP mice (The Jackson Laboratory; catalog no. 003946) to remove the NeoR cassette. Tail DNA from mice were PCR amplified and sequenced to confirm the desired truncation. Genotyping was performed with primers (5′-CTGGCACTGACCACTTGCTA-3′ and 5′-AACCGCTGCTCATTCTCTGT-3′) flanking the flippase recognition target site left after NeoR deletion. All animal work was conducted in a pathogen-free facility, and all the procedures were approved by the Institutional Animal Care and Use Committee at Columbia University Medical Center.

### Immunofluorescence

U2OS cells were seeded on coverslips 24 hours before transfection. Staining was done after another 18∼24 hours. Cells were washed with warm 1x PBS and fixed with 4% paraformaldehyde in PBS for 25 minutes at room temperature. Before staining, the cells were permeabilized with 0.1% triton X-100/PBS for 10 minutes and blocked with 3% BSA for 1 hour at room temperature. Fixed cells were then incubated with primary antibody anti-DDX21 (Novus, NB100-1718, 1:500) in 3% BSA for 1 hour at room temperature, followed by fluorophore-conjugated secondary antibody (Alexa Fluor 594-conjugated anti-rabbit, Invitrogen, 1:500) for 1 hour at room temperature. All images were captured on Nikon A1RMP confocal microscope with a 63× objective.

### Live-cell Imaging data collection and processing

Immortalized mouse embryonic fibroblast (iMEF) cells were plated in 35 mm diameter glass-bottom dishes. On the next day, the cells were transfected with plasmids encoding GFP-tagged mouse Ku70^WT^ or Ku70^ΔSAP^ via Lipofectamine 2000 (Invitrogen, Cat. 11668019), if indicated, the cells were treated with 20 µM of 5-bromo-2’-deoxyuridine (BrdU, Sigma) at 4 hours after transfection. Live-cell imaging was performed 24 hours after transfection with a Nikon Ti Eclipse inverted microscope equipped with the A1RMP confocal microscope system and Lu-N3 Laser Units. Laser micro-irradiation and timelapse imaging were carried out using the NIS Element High Content Analysis software and a 405 nm laser (energy level ∼500 uW for a ∼0.8 μm diameter region). Images were acquired every 10 s after micro-irradiation for a total of 5 min and then acquired every 5 min until 30 min after micro-irradiation. The relative intensity at damaged sites was calculated as the ratio of the mean intensity at each micro-irradiation damaged site to the corresponding mean intensity of the nucleus as background. Fiji software was used for quantifications and at least eight individual cells were analyzed for each data point in each experiment. At least two independent experiments were conducted.

### Lymphocyte development analyses

Single-cell suspensions were prepared from bone marrow, spleen and thymus of young adult (8–12 weeks) mice. Splenocytes were treated with red blood cell lysis buffer (Lonza ACK Lysis Buffer) for 4–5 min at room temperature to remove enucleated erythrocytes before staining. Approximately 2.5 × 10^6^ cells were stained using fluorescence-conjugated antibodies and analyzed by flow cytometry. The following antibody mixtures were used for B cell (FITC rat anti-mouse CD43, BD Pharmingen 553270; PE goat anti-mouse IgM, Southern Biotech 1020-09; PE-Cy5 anti-Hu/Mo CD45R (B220), eBioscience 15-0452-83; and APC anti-mouse TER-119, BioLegend 116212) and T cell (FITC anti-mouse CD8α, BioLegend 100706; PE rat anti-mouse CD4, BD Pharmingen 557308; PE/Cy5 anti-mouse TER-119, eBioscience 15-5921-82; and APC hamster anti-mouse TCRβ, BD Pharmingen 553174) analyses. The dead cells and debris were excluded based on their high side scatter and low forward scatter. Ter119 (an early proerythroblast to mature erythrocyte marker) antibody was used to gate out the erythroid cells from early proerythroblast to mature erythrocyte stages before the analyses of B and T cell-specific markers as in Fig. 2A. Flow cytometry was performed on an Attune flow cytometer (Thermo Fisher), and data was processed using FlowJo.

### Chromosomal V(D)J recombination assay

Total bone marrow was isolated from 4-week-old mice with Eμ-Bcl2 transgene and infected with retrovirus encoding v-abl kinase (75). Cells were cultured in DMEM with 15% fetal bovine serum for at least two months to achieve stable transformation. Chromosomal V(D)J recombination substrate (pMX-INV) was integrated into the cell lines through retrovirus infection, followed by positive selection via magnetic bead purification. V(D)J recombination with integrated substrate was carried out as before (67,75). Briefly, v-abl transformed B cells with stable integration of the substrate were treated with STI571 (3 μM, Novartis Pharmaceuticals) for 2 and 3 days. FACS was run for GFP expression and genomic DNA was collected for Southern blot. Specifically, DNA from individual lines was prepared on day 0 (before treatment), day 2, and day 3 after STI571 treatment, digested with with either EcoRV or EcoRV + NcoI, and Southern blotted with the GFP or C4 probe probe (Fig. 3G). The GFP probe (399 bp) was obtained by PCR amplification using primers (5′-AAGGACGACGGCAACTACAAG-3′, and 5′-CATGCCGAGAGTGATCCC-3′) with pMX-INV plasmid as template. The C4 probe is the 982 bp fragment generated by digestion with HindIII and NheI of the pMX-INV substrate plasmid.

### Class switch recombination analysis

The splenic CD43^−^ cells, isolated from 8–12 weeks old mice with CD43 magnetic beads (MACS, Miltenyi Biotec), were cultured in RPMI medium (Gibco) supplemented with 15% FBS, 1 μg/mL anti-CD40 (BD Pharmingen, 553721) and 20 ng/mL IL-4 (R&D, 404-ML-050) at a density of 1 × 10^6^ cells/mL. Cells were analyzed by flow cytometry after staining for IgG1 (FITC anti-IgG1, BD Pharmingen 553443) and B220 (PE/Cy7 anti-B220, BioLegend 103222) on days 3 and 4.

### High-throughput genomic translocation sequence (HTGT-seq)

HTGT-seq was performed as described (55,76,77). To study the junctions from chromosomal V(D)J recombination, 20 μg genomic DNA from v-abl transformed B cells with stable integration of the pMX-INV substrate was sonicated. The biotin-labeled primer (/5Bios/GGCAGCCTACCAAGAACAAC) was used for the linear amplification and another primer (5′-CGGTGGTACCTCACCCTTA-3′) was applied for nested amplification. Germline (unrearranged) sequence was removed with EcoRI (New England Biolabs) digestion. For the sequence analysis, the pMX-INV substrate sequence was added as an additional chromosome on top of the mm10 genome during the alignment process. To study the junctions from class switch recombination, 20 μg genomic DNA from CD43^−^ B cells stimulated with anti-CD40 and IL-4 for 4 days, was sonicated and amplified with an Sμ-specific biotin primer (/5Bios/CAGACCTGGGAATGTATGGT) and nested primer (5′-CACACAAAGACTCTGGACCTC-3′). AflII (New England Biolabs) was used to remove germline sequence. Since all our experimental mice and the pre-assembled IgH were of pure 129 background, the IgH switch region (from JH4 to the last Cα exon, chr12 114494415–114666816) of the C57/BL6-based mm9 genome was replaced with the corresponding region from the AJ851868.3 (GenBank accession no. AJ851868.3) 129 IgH sequence (1415966–1592715) to generate the mm9sr (switch region replacement) genome. Sequences were analyzed as detailed before (55,76) with the pipeline deposited on GitHub (https://github.com/robinmeyers/transloc_pipeline). Best-path searching algorithm (related to YAHA (78)) was used to select optimal junctions from Bowtie2-reported top alignments (alignment score > 50). The reads were then filtered to exclude misprimed events, germline sequence, competing prey and duplicated reads. A “duplicate read” was defined by the coordinates of a bait and prey alignment within 2 nt of another bait and prey alignment. Since the signal joint reads number was very low after applying the deduplication filter, this filter was not used for SJ analysis. SJ deletion size was capped at 1,000 bp. Since some switch region sequences are very repetitive, the alignments in those regions are filtered out by the mappability filter. But they do unequivocally map to an individual switch region. To plot all the switch region junctions, we added the ones filtered out by the mappability filter back after deduplication (76). MHs are defined as regions of 100% homology between the bait and prey-break site. Insertions are defined as regions containing nucleotides that map to neither the bait nor prey-break site. Blunt junctions are considered to have no MHs or insertions.

### Alphafold 3 modeling and ChimeraX

The full length of human Ku70 and the core of human Ku80 (1-545aa) were used to generate all the models using the Alphafold 3 sever (https://alphafoldserver.com/about). We removed the flexible linker, C-terminal region and the tail from the Ku80, since they were largely randomly placed within the top 5 predictions likely due to the long flexible linker. 25 bp dsDNA and corresponding dsRNA were also included. (dsDNA: 5’-<u>GTTTTTAGTTTATTAGCTTAGCTCCG</u>ATTT<u>CGGAGCAAGCTAATAAACTAAAAAC</u>-3’ (the underlined are the complementary bases) The corresponding dsRNA sequence is 5’-<u>GUUUUUAGUUUAUUAGCUUAGCUCCG</u>AUUU<u>CGGAGCAAGCUAAUAAACUAAAAAC</u>-3’.) The results were displayed using UCSF ChimeraX (https://www.cgl.ucsf.edu/chimerax/). All top 5 predictions were overlayed and colored by chain. B-factors, also known as temperature factors, are values that quantify the uncertainty of each atom in a protein crystal. The B-factor of the top prediction is displayed via ChimeraX. The red is high and blue is low.

### Surface Plasmon Resonance (SPR) Assay

SPR assays were implemented on Biacore T200 (Cytiva) using a buffer (20 mM Tris-HCl (7.5), 150 mM KCl, 1 mM DTT, 0.02% CA-630) at 25°C. A DNA substrate of 400 bp with a biotin at one end was prepared by PCR reaction on PhiX174 DNA, and was injected on a Series S Sensor Chip SA (GE Healthcare) installed in Biacore T200. Indicated concentrations of KU WT or ΔSAP mutant were applied to the sensor chip with DNA described above, in the following settings: contact time, 180 s; dissociation time, 300 s; flow rate, 30 μl/min. SPR data were analyzed on software installed in Biacore T200.

### DNA construct preparation for single-molecule colocalization

HPLC-purified oligonucleotides were obtained from Integrated DNA Technologies (IDT, Coralville, IA). To prepare a 10 µM solution of a 30 bp DNA construct, the following oligonucleotides were used: oligonucleotide 1 (5’ /Biosg/CCT AAT CTC ACA TGG CGA CGG CAG CGA GGC) and oligonucleotide 2 (5’ GCC /iCy3N/CG CTG CCG TCG CCA TGT GAG ATT AGG). These oligonucleotides were annealed to form the DNA construct. The resulting construct was stored at -20°C.

### Fluorescent labelling of Ku

Ku-SNAP tag protein and SNAP-Surface Alexa Fluor 647 (New England Biolabs, NEB) was mixed and incubated at 4°C for 3 hours. Following incubation, the mixture was subjected to five wash cycles using wash buffer (20 mM Tris, 150 mM KCl, 1 mM DTT, 1 mM EDTA, pH 7.4). Washing was performed using an Amicon filter with a 30K MWCO (Millipore) at 7,000 × g to remove unbound Alexa Fluor 647. After purification, glycerol was added to the protein solution to achieve a final concentration of 10% glycerol. The labelling efficiency of the Ku(wt) SNAP, and Ku(delSAP) SNAP proteins was assessed to be 49.4% and 52.5% respectively using spectrophotometric analysis. The labelled protein was aliquoted and stored at -80°C.

### Flow channel preparation for single-molecule colocalization experiment

Coverslips (Fisherfinest, 24×50 mm) and glass slides (Fisherfinest, 25×75×1 mm) were first cleaned by immersing in 4 M KOH (Cat. 06005-1KG, Honeywell) for 30 minutes. Following this, they were thoroughly washed with distilled water and methanol. To functionalize the glass surfaces with primary amine groups, a mixture of methanol (LC168104, Labchem), glacial acetic acid (BP1185, Thermo Fisher Scientific), and 3-(2-Aminoethylamino)propyltrimethoxysilane (Cat. A0774, TCI) in a ratio of 100:2:5 was applied. Glass slides and coverslips were washed, dried, and incubated with mPEG-SVA-5000 (Laysan Bio) and a mixture of mPEG-SVA-5000 and Biotin-PEG-SVA-5000 (Laysan Bio), respectively overnight in the dark. After incubation, the coverslips and glass slides were washed with distilled water, dried, and stored at -20°C. For microfluidic flow channel assembly, double-sided tape was used to sandwich between the pegylated glass slide and the biotin-functionalized coverslip. Epoxy was applied to seal the open ends of the flow channel. Neutravidin solution (10 µL, Invitrogen, Life Technologies Corporation, Eugene, OR) was injected through the drilled holes in the glass slide and incubated for 10 minutes. Excess neutravidin was removed by washing with 50 µL of BSA solution (30 mg/mL, Gemini Bio-products), followed by a 10-minute incubation to block any remaining reactive sites.

### Total internal reflection fluorescence (TIRF) microscope setup

The experiment was performed using a Total Internal Reflection Fluorescence (TIRF) microscope, which featured an inverted oil immersion objective with a 100x magnification and a 1.49 numerical aperture. The setup included 532 nm and 639 nm excitation lasers (Ultra-Laser, MGL-FN-532 and MGL-FN-639, respectively) for TIRF illumination. A dichroic beam splitter (Semrock, Di03-635-t1) separated the green and red fluorescence signals, which were then filtered through narrow bandpass filters (Semrock, FF01-582/64 and FF01-680/42). These signals were captured by an electron-multiplying charge-coupled device (EM-CCD) camera (Andor, iXon897). Fluorescence beads (Invitrogen, Life Technologies Corporation, Eugene, OR) were utilized to calibrate the green and red signal channels.

### Single-molecule colocalization experiment

The flow channel was first washed with 20 mM Tris buffer containing 100 mM KCl and 2.5 mM MgCl₂. Following this, 35 μL of a 50 pM 30 bp dsDNA construct was injected into the microfluidic channel, incubated for 3 minutes, and then washed with the same buffer. Next, an imaging buffer composed of 20 mM Tris, 100 mM KCl, 2.5 mM MgCl₂, 1 mg/mL BSA, 0.8% glucose, 0.1% Trolox, 2 mM DTT (adjusted to pH 7.5), and 1% GLOX was introduced into the channel containing unlabelled Ku protein. This mixture was incubated for 5 minutes. Afterward, labelled Ku protein was injected, and imaging was performed using a TIRF microscope at various time intervals.

For the exchange assay, the incubation with unlabelled Ku protein was replaced with the same protein in its labelled form (e.g., unlabelled Ku(wt) flushed with labelled Ku(wt)-AF647 (Fig S9B top), and preincubated unlabelled Ku(delSAP) flushed with labelled Ku(delSAP)-AF647 (Fig S9B bottom)).

For the displacement assay, the unlabelled Ku protein was replaced with an alternative labelled protein (e.g., unlabelled Ku(delSAP) flushed with labelled Ku(wt)-AF647 (Fig S9C top), and preincubated unlabelled Ku(wt) flushed with labelled Ku(delSAP)-AF647 (Fig S9B bottom)).

Movies of 1000 frames at 33 Hz were recorded, with red lasers active for the first 900 frames and both lasers on for the final 100 frames. Three independent experiments were conducted. The percentage of colocalization between the stable red signal (representing the Ku molecule) on the green DNA signal was calculated and corrected for the labelling efficiency of the respective proteins.

## Supplementary Data Figure

**Figure S1.**
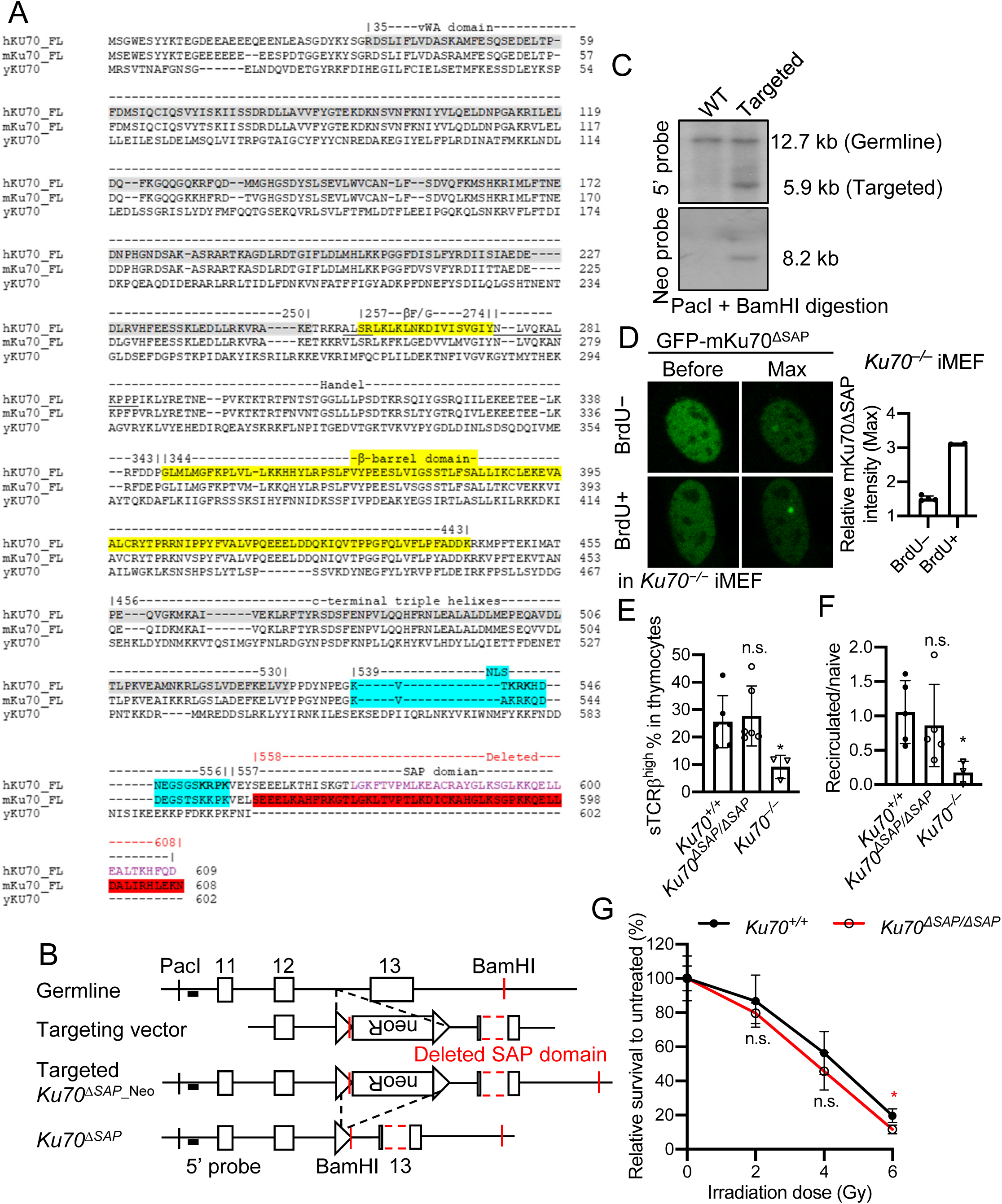
Generation of *Ku70^ΔSAP^* allele and mouse models. (A) The alignment of Ku70 protein sequences from human, mouse and yeast. The beta-barral domain is highlighted in yellow, the nuclear localization signal (NLS) in turquoise, and the SAP domain (deleted) in red. (B) The targeting scheme of *Ku70^ΔSAP^* allele. Murine Ku70 locus (top), targeting vector (second row), targeted allele (third row), and the neo-deleted allele (bottom). The exons and flippase recognition target sites are shown as open boxes and triangles, respectively. All exons on the arms are marked. The 5′ probe is marked as a thick black line. The recognition sites of PacI and BamHI used for southern blot are marked. The red dashed box in exon 13 represents the locus of the SAP domain we deleted. (C) Southern blot analyses of PacI and BamHI digested DNA from *Ku70^+/+^* and *Ku70^ΔSAP^* targeted ES cells, blotted with the 5′ or Neo probe. (D) Representative images of GFP-KU foci with or without BrdU sensitatation. (E) The percentage of surface-TCRβ^high^ (the right-side peak in Fig. 2*A*) cells in thymus. (F) Quantification of recirculated B-cell (B220^high^IgM^+^) versus naïve B-cell (B220^mid^IgM^+^) ratio in the bone marrow. For panels *D* and *F*, the bars represent the average and the standard deviation of 3 or more mice per genotype. Student’s *t* test was used to calculate the *P* value. n.s.: no significant difference. *: *P* < 0.05. (*G*) The sensitivity to irradiation (IR) (2, 4 and 6 gray (Gy)) of *Ku70^+/+^* and *Ku70^ΔSAP/ΔSAP^* B-cells activated for CSR. The average ± standard deviation from each IR dose were plotted. Student’s *t* test was used to calculate the *P* value of each IR dose. n.s.: no significant difference. *: *P* < 0.05. Triplicates of each genotype were assayed in two independent experiments.

**Figure S2.**
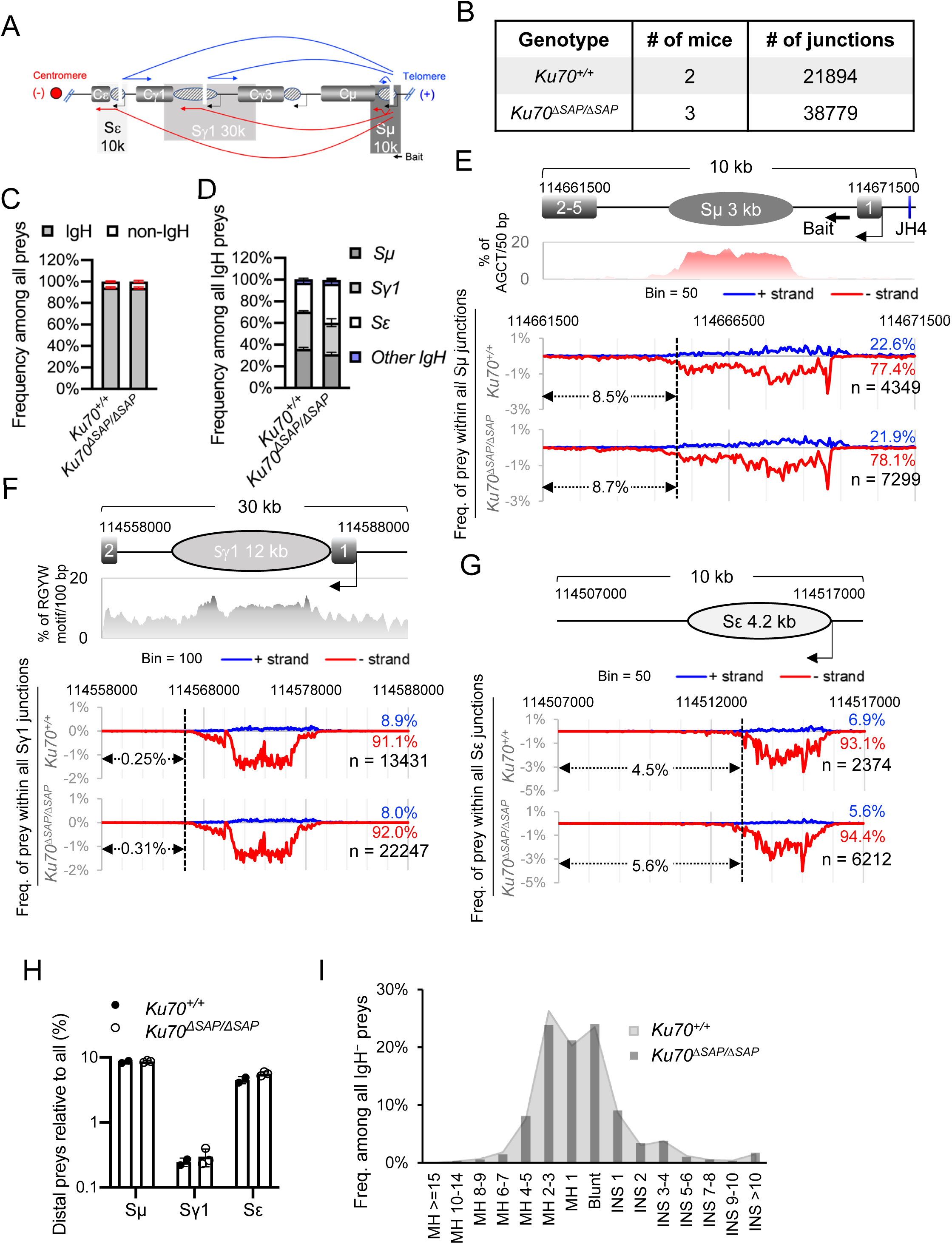
HTGT-seq analyses of CSR to Sμ, Sγ1 and Sε region using an Sμ bait. (A) Diagram of the murine IgH locus with the location of Sμ (dark grey), Sγ1 (grey), and Sε (light grey). The bait site at the 5′Sμ region is also marked. Since IgH resides on the minus strand of murine chromosome 12, the preys on the (−) strand (red arrows) reflect normal internal deletion and CSR, and the preys on the (+) strand (blue arrows) indicate inversion or inter-chromosomal junction with a sister or homologous chromosome. (B) Summary of the CSR libraries and junctions analyzed by HTGT-seq. (C) The percentage of IgH and non-IgH junctions among all junctions in each genotype. (D) The relative frequency of junctions in Sμ, Sγ1, Sε and other regions of IgH locus among all junctions by genotype. (E) The spatial distribution of preys in Sμ region. Schematic of the Sμ region with the bait site at the 5′ Sμ marked by an arrow is at Top. The frequency of AGCT motifs per 50-bp double-strand DNA is plotted. The values between y-axis and the dashed line (114,661,500∼114,665,100) indicate the percentage of preys that fall outside the core Sμ region. (F) The spatial distribution of preys in Sγ1 region. Schematic of the Sγ1 region is at Top. The frequency of RGYW motifs per 100-bp double-strand DNA is plotted. The values between y-axis and the dashed line (114,558,000∼114,566,000) indicate the percentage of preys that fall outside the core Sγ1 region. (G) The spatial distribution of preys within Sε region (bin size = 50 bp). Schematic of the Sε region is at Top. The values between y-axis and the dashed line (114,507,000∼114,513,000) indicate the percentage of preys that fall outside the core Sε region. For panels *E–G*, all (−, red, below zero) and (+, blue, above zero) strand preys add up to 100%. For each genotype, the percentage of (+) strand (blue) and (−) strand (red) junctions and total number are marked at the right. The data from multiple mice of the same genotype was pooled and plotted together (see panel *B* for sample information). (H) Relative frequency of distal preys among the preys in Sμ, Sγ1 and Sε region. (I) The distribution of all IgH junctions by junctional type (MH, microhomology. INS, insertion). This graph represents the pool of at least two independent libraries (see panel *B* for sample information) of each genotype.

**Figure S3.**
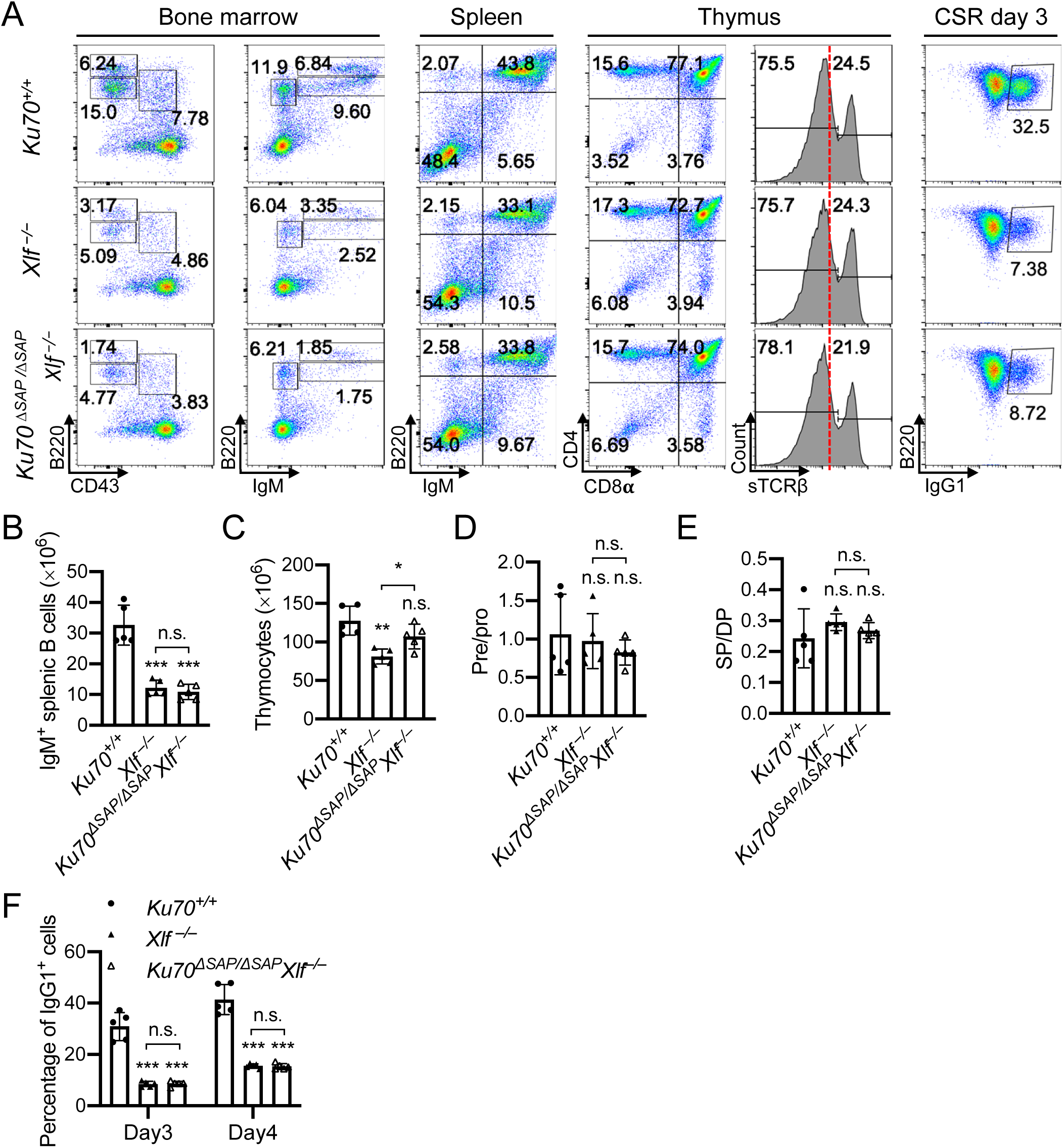
*Ku70^ΔSAP/ΔSAP^Xlf ^−/−^* mice have normal lymphocyte development and maturation. (A) Representative flow cytometric analyses of bone marrow, spleen and thymus from *Ku70^+/+^*, *Xlf ^−/−^* and *Ku70^ΔSAP/ΔSAP^Xlf ^−/−^* mice. Numbers on the plot are percentages of cells among the gated populations. (B and C) The total IgM^+^ splenic B-cell numbers (B) and total thymocyte numbers (C). (D) The ratio of pre-B (B220^+^IgM^−^CD43^−^) versus pro-B (B220^+^IgM^−^CD43^+^) cells in the bone marrow. (E) The ratio between the sum of CD4^+^ or CD8^+^ single positive T-cells (SP) and CD4^+^CD8^+^ double positive (DP) immature T-cells in the thymus from mice of different genotypes. (F) Quantification of B220^+^IgG1^+^ B-cell percentage on day 3 and 4 of stimulation. For panels *B–F*, the bars represent the average and the standard deviation of 5 mice per genotype. Student’s *t* test was used to calculate the *P* value. n.s.: no significant difference. *: *P* < 0.05, **: *P* < 0.01, ***: *P <* 0.001.

**Figure S4.**
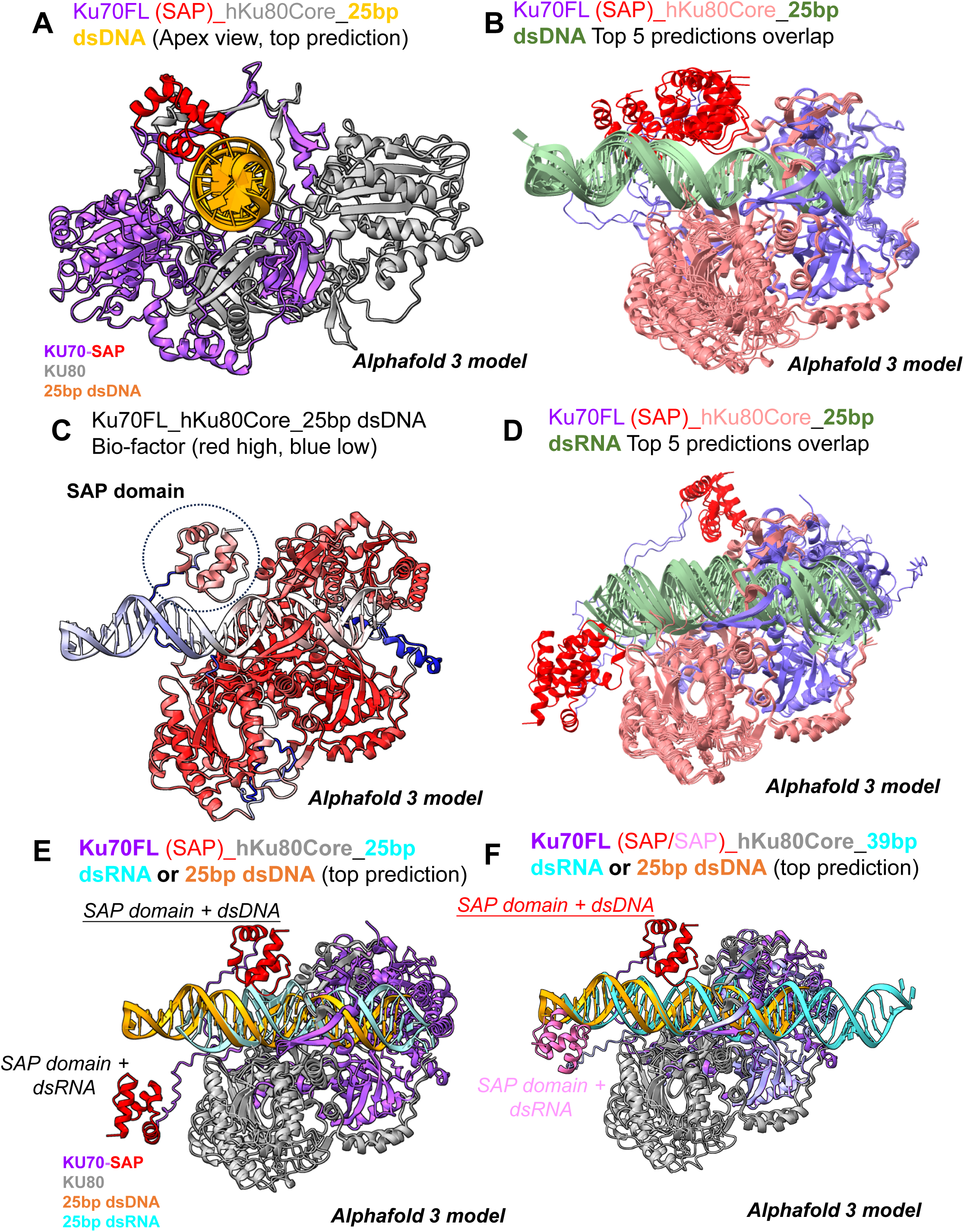
Alphafold 3 model of human KU complex on 25bp dsDNA and dsRNA. (A) The AlphaFold 3-generated structural model (top prediction) of the full-length human KU70 (purple with SAP domain in red), the core region of human KU80 (1-545 aa, grey), and a 25 bp dsDNA (bright orange). (B) Overlays of the top five predictions with 25 bp dsDNA. KU70 is colored pink with the SAP domain in red, and the core of KU80 is in purple. (C) B-factor display of the top prediction on 25 bp dsDNA, where red indicates high stability and blue indicates low stability. (D) Overlays of the top five predictions with 25 bp dsRNA (same sequence as the dsDNA). KU70 is colored pink with the SAP domain in red, and the core of KU80 is in purple. Unlike the dsDNA model, the SAP domain is not consistently placed when dsRNA is used. (E) Overlay of the top prediction with 25 bp dsDNA (bright orange) and 25 bp dsRNA (turquoise).(F) Overlay of the top prediction with 25 bp dsDNA (bright orange) and 39 bp dsRNA (turquoise). The SAP domain from the dsRNA complex is shaded in pink.

**Figure S5.**
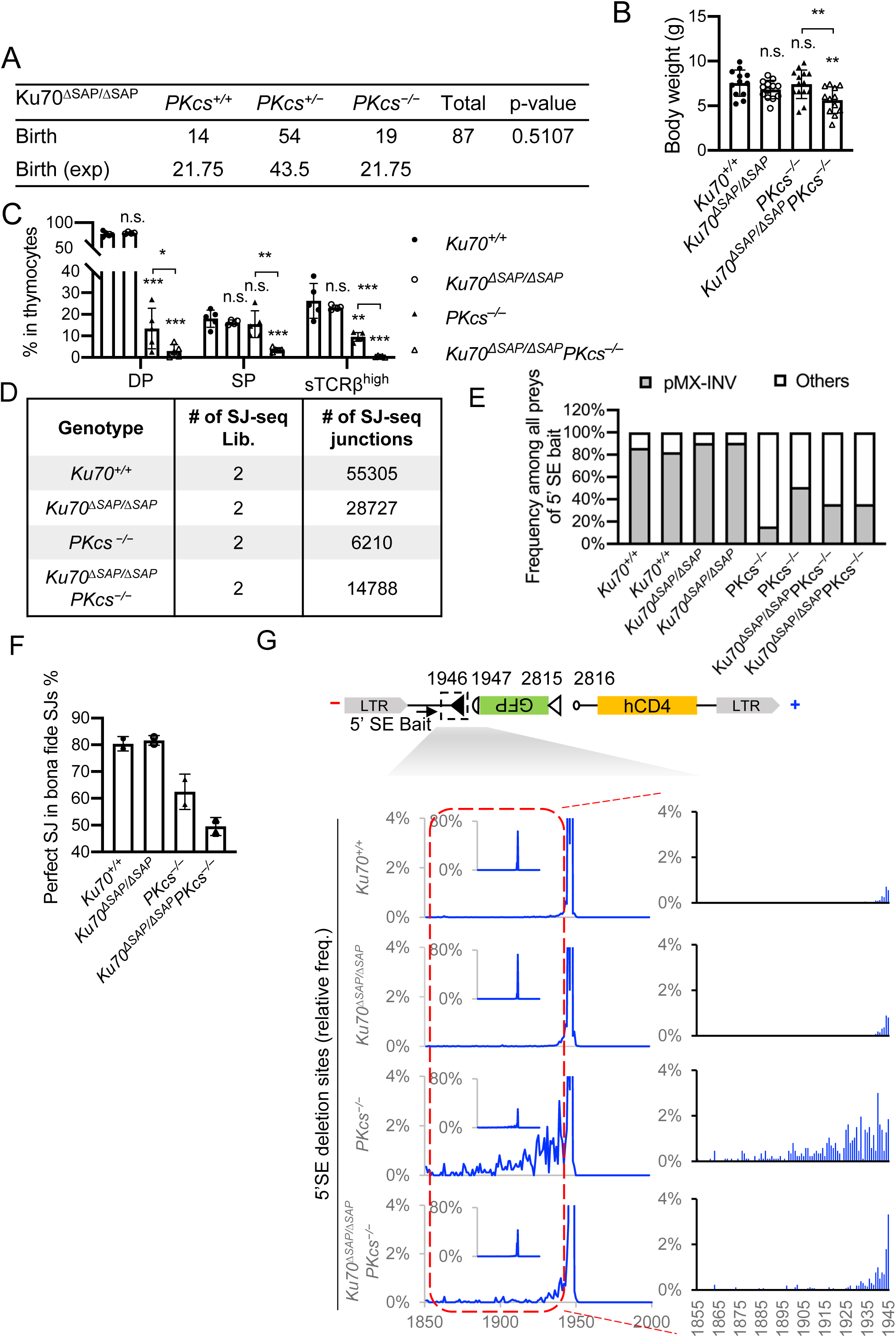
SAP domain of Ku70 is important for lymphocyte development and V(D)J recombination in *DNA-PKcs^−/−^* mice. (A) The number of live born mice obtained at P7 from crosses between *DNA-PKcs^+/−^Ku70^ΔSAP/ΔSAP^* mice. The *P* value was calculated with the chi-squared test. (B) The total body weight of mice (P7–P10) from the different genotypes indicated. (C) The percentage of CD4^+^CD8^+^ DP immature T-cells, the sum of CD4^+^ or CD8^+^ SP T-cells and surface-TCRβ^+^ (the right-side peak in Fig. 3*A*) cells in total thymocytes. For panels *B* and *C*, the bars represent the average and the standard deviation of at least 4 mice per genotype. Student’s *t* test was used to calculate the *P* value. n.s.: no significant difference. *: *P* < 0.05, **: *P* < 0.01, ***: *P <* 0.001. (D) Summary of the libraries and junctions analyzed by HTGT-seq by genotype. The total number of junctions are listed. (E) Frequency of recovered HTGT-seq junctions amplified by 5′SE bait primer mapped to the pMX-INV substrate or elsewhere in the genome (others) from two independent repeats of G1 arrested samples. (F) The frequency of precise SJ among all bona fide SJs recovered in each genotype. The bars represent the average and standard deviation of two samples per genotype. (G) (Top) Schematic diagram of pMX-INV substrate for V(D)J recombination with the location of the 5′ bait SE RSSs (the filled black triangle). (Bottom) The spatial distribution of 5′SE terminal locations among all junctions within pMX-INV substrate (1 bp bin).

**Figure S6.**
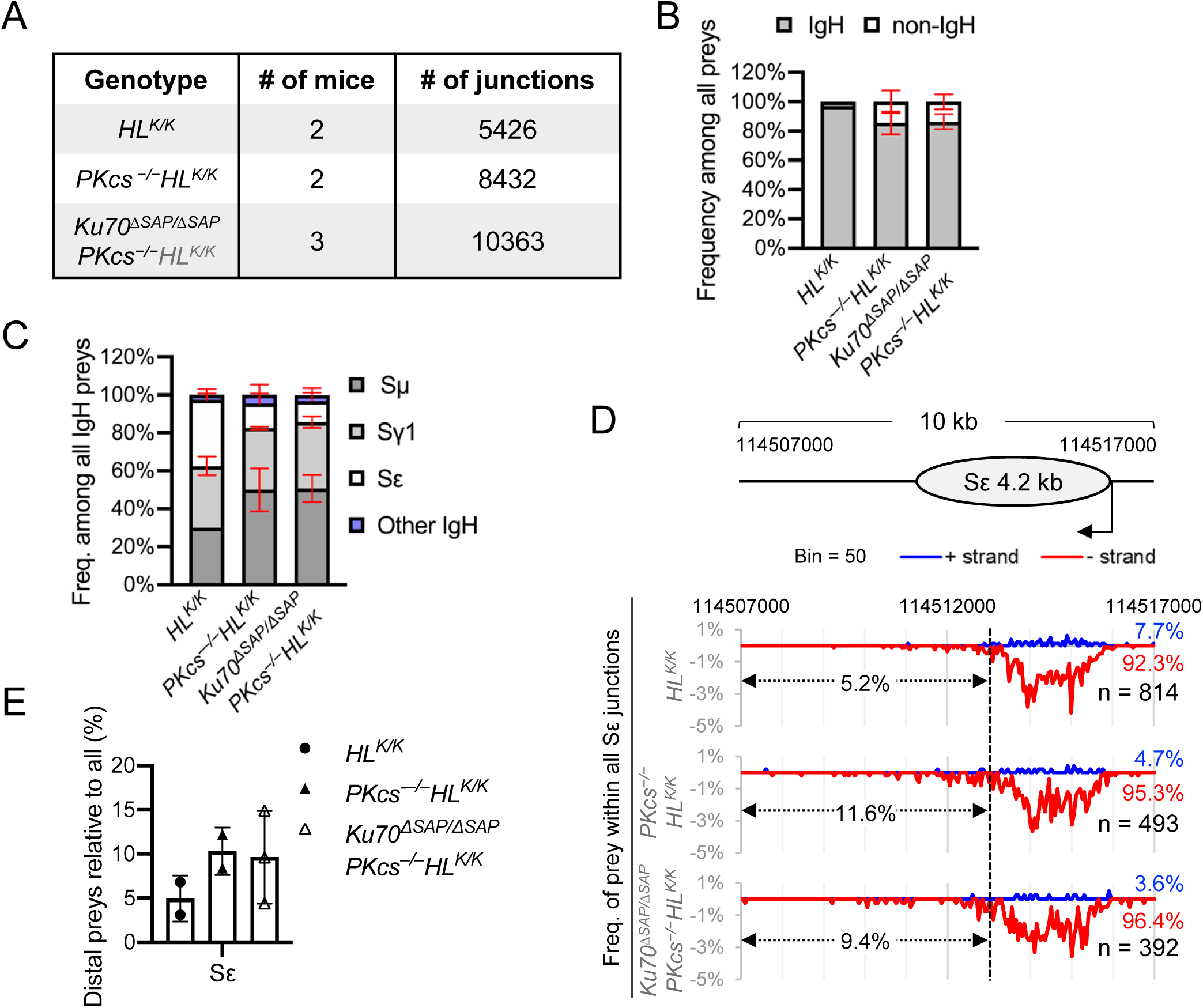
HTGT-seq analyses of CSR to Sε region using an Sμ bait. (A) Summary of the CSR libraries and junctions analyzed by HTGT-seq. (B) The percentage of IgH and non-IgH junctions among all junctions in each genotype. (C) The relative frequency of junctions in Sμ, Sγ1, Sε and other regions of the IgH locus among all junctions within the IgH by genotype. (D) The spatial distribution of preys within the Sε region (bin size = 50 bp). The data from multiple mice of the same genotype was pooled and plotted together (see panel *A* for sample information). Schematic of the Sε region is at Top. All (−, red, below zero) and (+, blue, above zero) strand preys add up to 100%. For each genotype, the percentage of (+) strand (blue) and (−) strand (red) junctions and total number are marked at the right. The values between y-axis and the dashed line (114,507,000∼114,513,000) indicate the percentage of preys that fall outside the core Sε region. (E) Relative frequency of distal preys among the preys in Sε region.

**Figure S7.**
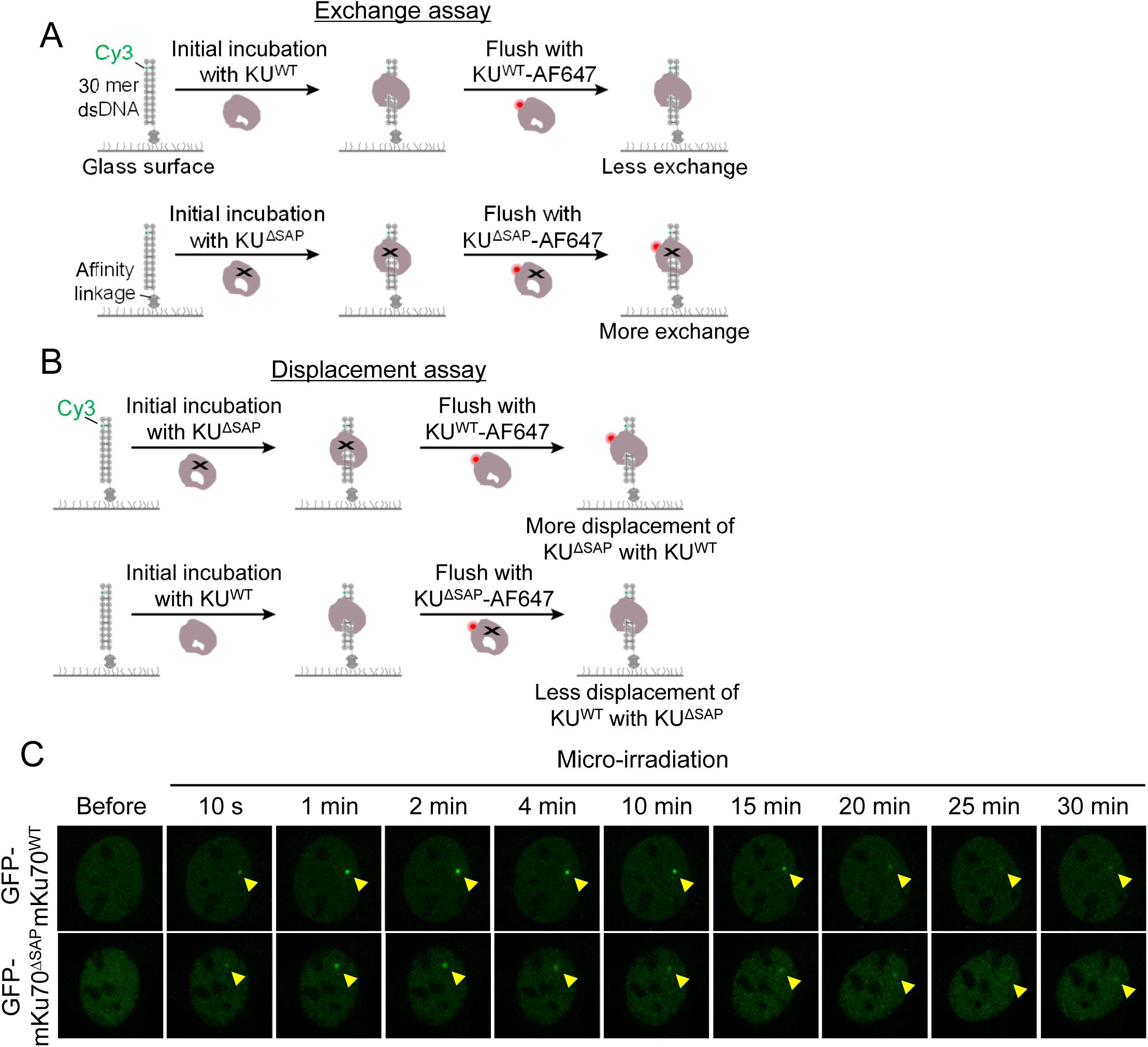
Ku70-SAP domain is critical for KU loading *in vitro*. (*A*) Schematic representation of exchange assay. Cy3 labeled 30mer DNA incubated with KU^WT^ (top) and KU^ΔSAP^ (bottom) for 5 min followed by flushing with the same protein labeled with AF647. (B) Schematic representation of displacement assay. (Top) Cy3-labeled 30mer DNA incubated with KU^ΔSAP^ for 5min followed by flushing with AF647-labeled KU^WT^. (Bottom) Cy3-labeled 30mer DNA incubated with KU^WT^ for 5min followed by flushing with AF647-labeled KU^ΔSAP^. (C) Representative images of micro-irradiation-induced GFP-mKu70^WT^ or GFP-mKu70^ΔSAP^ foci in Ku70 knockout iMEF cells. The yellow arrowheads point to the area of micro-irradiation.

